# Design and deployment of a regulation-compliant infrared heating system for UK field trials

**DOI:** 10.64898/2026.01.19.700297

**Authors:** Isabel Faci, Paul Rogerson, James Simmonds, Martyn Hewitt, Darryl Playford, Antony N. Dodd, Cristóbal Uauy

## Abstract

Free-air warming systems are essential for simulating climate change in field conditions, yet most are not designed to meet UK or EU health and safety standards. We present the design and field deployment of a hexagonal Temperature-Free Air Controlled Enhancement (T-FACE) system that fully complies with these regulations. The system uses real-time, temperature-responsive control with multiple sensors to maintain a consistent temperature differential between heated and ambient temperature experimental plots. We describe its engineering design, performance, costs, and key operational challenges, particularly related to heater reliability under continuous use. We tested the T-FACE system during the winter months of a field sown wheat crop. Phenotypic evaluation showed effects of heating on heading date, plant height, spike length, and spikelet number, suggesting the system potential. This proof-of-concept provides a scalable, regulation-compliant approach for studying crop responses to climate warming under realistic field conditions.

## Introduction

Environmental cues such as temperature and photoperiod play a central role in regulating the timing of key developmental transitions in plants, including the transition to flowering. While controlled-environment studies enable dissection of the effects of individual cues, understanding how plants integrate multiple, fluctuating signals in natural conditions is equally important, particularly for predicting responses of crops under future climates. Field studies in *Arabidopsis thaliana* have underscored this need, showing that developmental outcomes can shift significantly under natural environmental variation, highlighting the limitations of controlled environments for predicting field performance (Hepworth et al., 2018, 2020; Wilczek et al., 2009, 2014). To address this, experimental systems like Temperature-Free Air Controlled Enhancement (T-FACE) systems enable the manipulation of environmental variables in the field, offering a valuable approach to study plant responses under varying climate scenarios.

In the UK, mean annual temperatures are projected to rise by approximately 3 °C by the 2061–2080 period under the *business-as-usual* Representative Concentration Pathway (RCP) 8.5 scenario (UKCP18, Met Office, 2023). A range of experimental approaches have been developed to simulate such future warming. For example, soil surface heating cables heat tissues near the ground but have little impact on canopy temperatures except in very short vegetation; however, when combined with mesh coverings, they can achieve more uniform heat distribution and affect canopy temperatures (Grime et al., 2000; Ineson et al., 1998; O’Neill et al., 2019). Nevertheless, soil heating combined with coverings can alter the microclimate, potentially influencing plant growth (Drost et al., 2017). Night-time warming methods, such as forced air mixing or insulating covers, have also been tested (Beier et al., 2004; Pinter, 2000), but these are typically only effective after sunset, offer limited control over timing, and cannot replicate daytime temperature dynamics.

Infrared (IR) heating as a free-air vegetation warming method, overcomes some of these limitations due to its ability to mimic solar radiation without the artefacts introduced by enclosures. The first IR system used parallel lines of heaters to apply radiant heat to plant canopies (Harte & Shaw, 1995). Later work showed that arranging heaters in hexagonal arrays angled at 45° produced more uniform warming (Kimball et al., 2008). This study also introduced feedback control using canopy temperature sensors to maintain a consistent warming differential relative to ambient conditions (Kimball et al., 2008). These systems have been deployed across various ecosystems and crop types, including grasslands with perennial ryegrass and white clover (Nijs et al., 1996), rice paddies (Zhang et al., 2020), and snow-covered subalpine forests (Meromy et al., 2015). Recent studies have shown their utility in capturing biologically relevant responses, such as reduced wheat (*Triticum aestivum*) nocturnal stomatal conductance and grain yield under night-time warming (McAusland et al., 2023).

These systems have been successfully deployed in North America, Mexico and East Asia, but to our knowledge, no published studies report their deployment in the UK or Europe. When exploring the potential to implement equivalent systems in the UK, we realised that these published designs were non-compliant with UK Health and Safety Executive (HSE) regulations, and were similarly non-compliant with relevant EU safety regulations. Here, we present the deployment of a hexagonal T-FACE warming system, specifically engineered to comply with all relevant UK HSE standards while maintaining a ∼ 3 °C warming differential consistent with 2060–2080 climate projections. We detail the system’s engineering design, control logic, performance, and deployment challenges, particularly those related to heater reliability during continuous operation. While the system successfully imposed sustained warming, technical limitations highlight the need for further refinement. This proof-of-concept study provides a valuable foundation towards enabling regulation-compliant, scalable field-based climate change experiments in UK cropping systems.

## Results

### Design and implementation of an UK-compliant free-air heating system with real-time temperature control

To simulate elevated temperatures under field conditions, we developed a hexagonal, free-air, infrared heating system with real-time temperature-responsive control (Figure 1). This design was based on that described by Kimball et al (Kimball et al., 2008), but with full compliance to UK HSE regulations. Each hexagonal array was equipped with six electric infrared heaters (2.6 kW, 3-10 µm wavelength) with Kanthal AF alloy heating elements and a surface temperature of up to 350 °C, enclosed in extruded aluminium. These heaters were UKCA and CE certified, meeting all relevant UK and EU safety standards for outdoor electrical equipment. The IP64 rating of these heaters ensures protection against dust and water ingress, and they emit no visible light, avoiding introducing Photosynthetically Active Radiation (PAR) as a confounding variable in the experimental set-up, while also preventing light disturbance to nearby residents. To further protect the heaters and prevent accidental contact, we installed custom aluminium covers (Figure 1a-b). We mounted the heaters at a 45° angle to create uniform heat distribution across the hexagonal plot (Figure S1a) (Kimball et al., 2008). To support the heaters, we used 48 mm aluminium key-clamp poles, secured in concrete footings for stability and weather resistance. All heaters were mounted on adjustable handrail brackets (Figure S1b) and positioned at 1.2 m above the canopy (Kimball et al., 2012).

**Figure 1.**
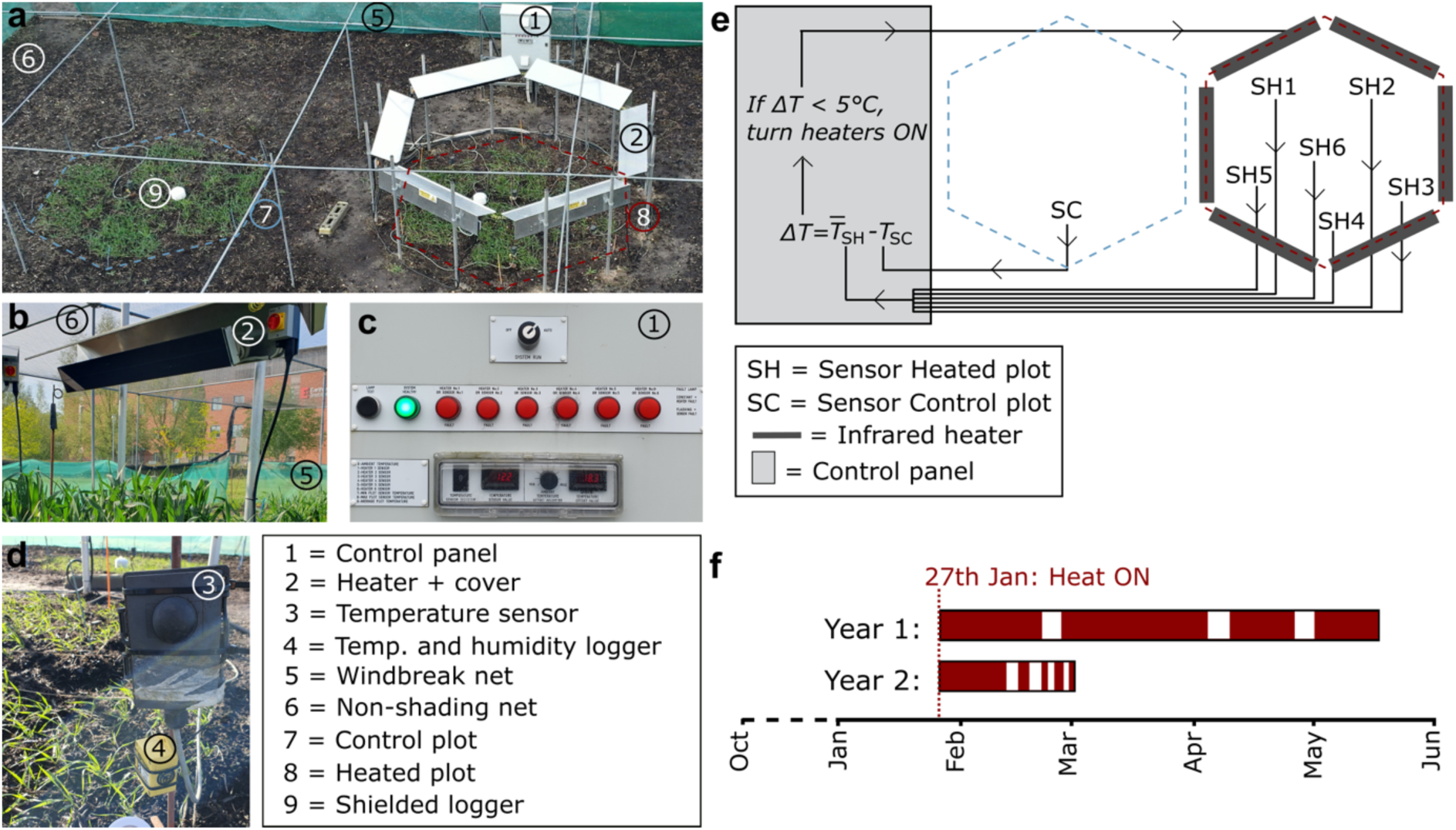
T-FACE system with real-time temperature control. **(a)** Aerial image of control (blue) and infrared heated (dark red) hexagonal plots. **(b)** Close view of black-body infrared-heater covered with aluminium cover, mounted on two aluminium poles. Next to the heater, rotary isolators (red switch in yellow squared panel). **(c)** Exterior view of the heating system control panel enclosure, with a key-operated switch, a lamp test button, and status indicators for system health (green) and faults (red) across six heaters. The bottom section provides temperature monitoring, displaying individual sensor readings (1–6), aggregate values (min/max/average), and an offset adjuster to select ΔT. **(d)** Close view of temperature sensor and temperature and humidity logger. Sensor elements of both are the visible circular black component, strategically positioned against direct sun. **(e)** Simple diagram of system logic; Temperature is measured in the control plot (SC) and at six locations in the heated plot (SH1–6). The control panel averages the heated plot readings, calculates the temperature difference (ΔT) relative to the control plot, and activates all six heaters when ΔT is below the target ΔT (+ 5 °C in this study). **(f)** Timing of heating applications across two seasons of an autumn-sown wheat crop: 2022–2023 (Year 1; 27^th^ January to 19^th^ May, 97 days effective heating) and 2023–2024 (Year 2; 27^th^ January to 5^th^ March, 25 days effective heating).

We next developed a control system to manage real-time temperature regulation. We supplied power from the mains electricity source to the control panel via a 32-amp, three-phase and neutral feed using 10 mm² multicore PVC/SWA/XLPE cable. From the panel, six 20-amp radial circuits (2.5 mm² PVC/SWA/XLPE cable) ran to the heaters, each terminating in a rotary isolator (20 A 4 Pole). Each circuit was protected by a Miniature Circuit Breaker (MCB), with a 40 A 30 mA residual current device (RCD) providing additional protection (Figure 1c, Figure S1c, Supplementary Material 1). To regulate heater output, we installed seven temperature sensors (Figure 1d): six inside the heated plot and one outside (within the control plot) as a reference. The control system continuously averaged the heated plot sensor readings and adjusted heating to maintain a fixed temperature difference above ambient (Figure 1e), which we set to + 5 °C using a selector switch on the control panel. Initial testing used fan speed controllers to modulate voltage and temperature, but these proved unreliable with prolonged use. We replaced them with solid-state relays, which were more reliable and simply turned all the heaters fully on or off as needed. The panel also displayed real-time temperatures from each sensor and system status, indicating whether it was operating normally or had stopped due to a fault (Figure 1c). To comply with UK HSE regulations, we enclosed all cables in weatherproof conduit, added warning signage to heater covers (Figure S1d), developed a standard operating procedure (SOP; Supplementary Material 2), and restricted site access with a locked perimeter (Figure S1e). The control panel was also sealed and protected in line with Institution of Engineering and Technology: Wiring Regulations BS 7671. These measures ensured continuous safe operation under UK HSE standards.

To monitor temperature and relative humidity (RH), we placed loggers inside and outside the heated plot, using both uncovered units (with sensors facing away from direct sun) and Stevenson screen–covered units. We compared the temperature differentials (ΔT) between heated and control plots for both logger types while the system was active. The mean ΔT from uncovered loggers was 1.6 °C higher than that from covered loggers (Wilcoxon test, *W* = 30,865, *p* < 2.2 × 10⁻¹⁶; Figure S1f). As each ΔT was calculated using a matching control (i.e. uncovered vs. uncovered; covered vs. covered), this discrepancy reflects the insulating effect of the Stevenson screens. Since the heating is primarily via infrared contact, the covers appear to block some of the applied heat, altering the recorded temperature. Therefore, all final temperature calculations were based on data from the uncovered loggers. To minimise heat loss from wind gusts across the heated plot, we installed windbreak netting around the perimeter of the plots, positioned > 2 m away from the crop. Additionally, we enclosed the sides and top with non-shading netting (also set back from the plants) to prevent animal entry and avoid crop damage (Figures 1a, 1b).

### T-FACE achieves between 2.3 °C to 5.6 °C temperature increase with relatively uniform heat distribution

We conducted proof-of-concept experiments across two seasons of an autumn-sown wheat crop: 2022–2023 (Year 1) and 2023–2024 (Year 2). Each year, we included a heated plot equipped with the T-FACE system and a control plot without heating infrastructure. Heating began in late January in both years, effectively running for approximately three months in Year 1 and nearly a month in Year 2 (Figure 1f).

To assess system performance, we examined the recordings from the uncovered temperature loggers inside and outside the heated plot. During 16 days of active heating in Year 2, the air temperature inside the plot was consistently higher that of the non-heated control plot (Figure 2a). Based on 30-minute intervals, 95% of ΔT values ranged from 2.3 °C to 5.6 °C (mean = 4.0 °C). Focusing on a single day temperature profile, the control plot shows a smoother temperature profile, while the heated plot exhibits a larger number of short, brief peaks which result from the heaters cycling on and off to maintain the target ΔT of + 5 °C.

**Figure 2.**
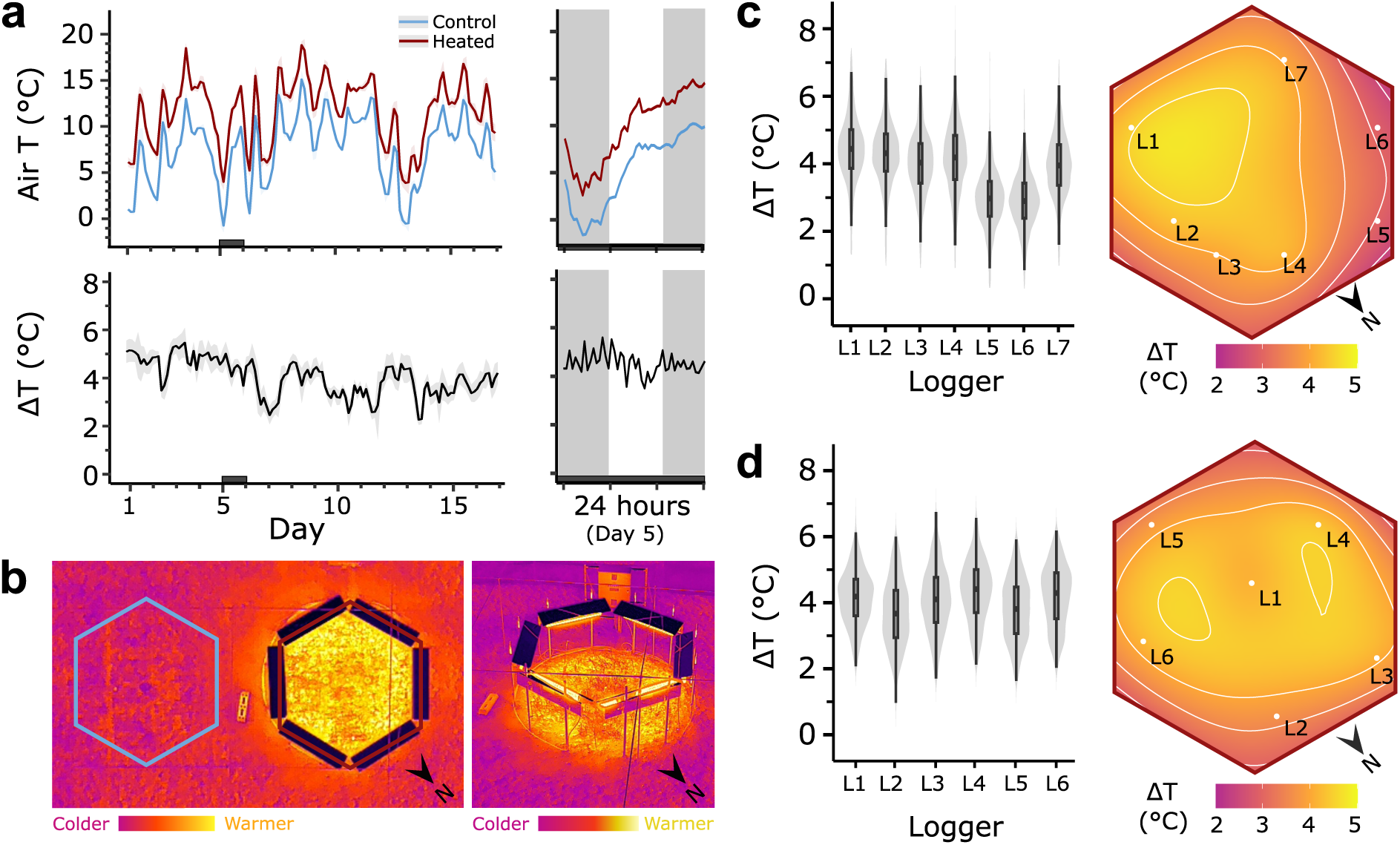
Differential temperature performance and spatial distribution of heating. **(a)** Air temperature (T) and differential temperature (ΔT = heated – control). Left, 16-day period when the heating system was continuously on (Year 2: 27 January to 11 February 2024). Each datapoint represents a 6h-window, averaged from 30-min intervals. The ribbon shading shows the variation within the 6h-window. Day 5 (31 January 2024) is highlighted with a dark rectangle and displayed at 30-min resolution on the right. Nighttime hours are shaded. **(b)** Drone thermal imaging pictures from Year 2 experiment (26 February 2024). Top view of control and heated plots (left) and oblique view (right) of heated plot. N indicates North. **(c)** Year 1, 36-day continuous heating period (1 March to 5 April 2023), 30-min windows: Left, distribution of ΔT recorded for each logger. Right, contour plot using the averaged ΔT for each logger across the heated plot. **(d)** Year 2, 16-day continuous heating period (27 January to 11 February 2024). Left, distribution of ΔT recorded for each logger. Right, contour plot using the averaged ΔT for each logger across the heated plot (see *Methods*).

To map heat distribution and explore heating boundaries, we obtained thermal images from above and at an angled top-side view. While qualitative, the images show relative uniform heat across the plot and a clear change in temperature at the edges, separating the heated hexagon from the surrounding area (Figure 2b). Spatial uniformity of heating was generally high, with individual sensors showing mean ΔT ranging from 3.67 °C to 4.44 °C and standard deviations from 0.83 °C to 1.06 °C, across both years (Figure 2c–d). Exceptions included two loggers in Year 1 (2.90 °C and 2.96 °C) located on the north-west side. Together, these results indicate relatively uniform heat distribution across the T-FACE plots.

### Assessing heating system performance

To further assess heater performance, we compared the ΔT between daytime and nighttime hours over a 16-day heating period in Year 1 and a 36-day heating period in Year 2. Day–night comparisons revealed slightly higher ΔT at night in both years (Wilcoxon test, Year 1: *W* = 322,615, *p-*adj < 0.001; Year 2: *W* = 61,725, *p-*adj = 0.027). In Year 1, mean ΔT was 3.70 °C during the day and 3.91 °C at night, while in Year 2 it was 3.93 °C during the day and 4.09 °C at night (Figure 3a). These differences, however, are relatively small in magnitude (ΔT = 0.21 °C and 0.16 °C, respectively) and indicate stable heater performance across day and night periods.

**Figure 3.**
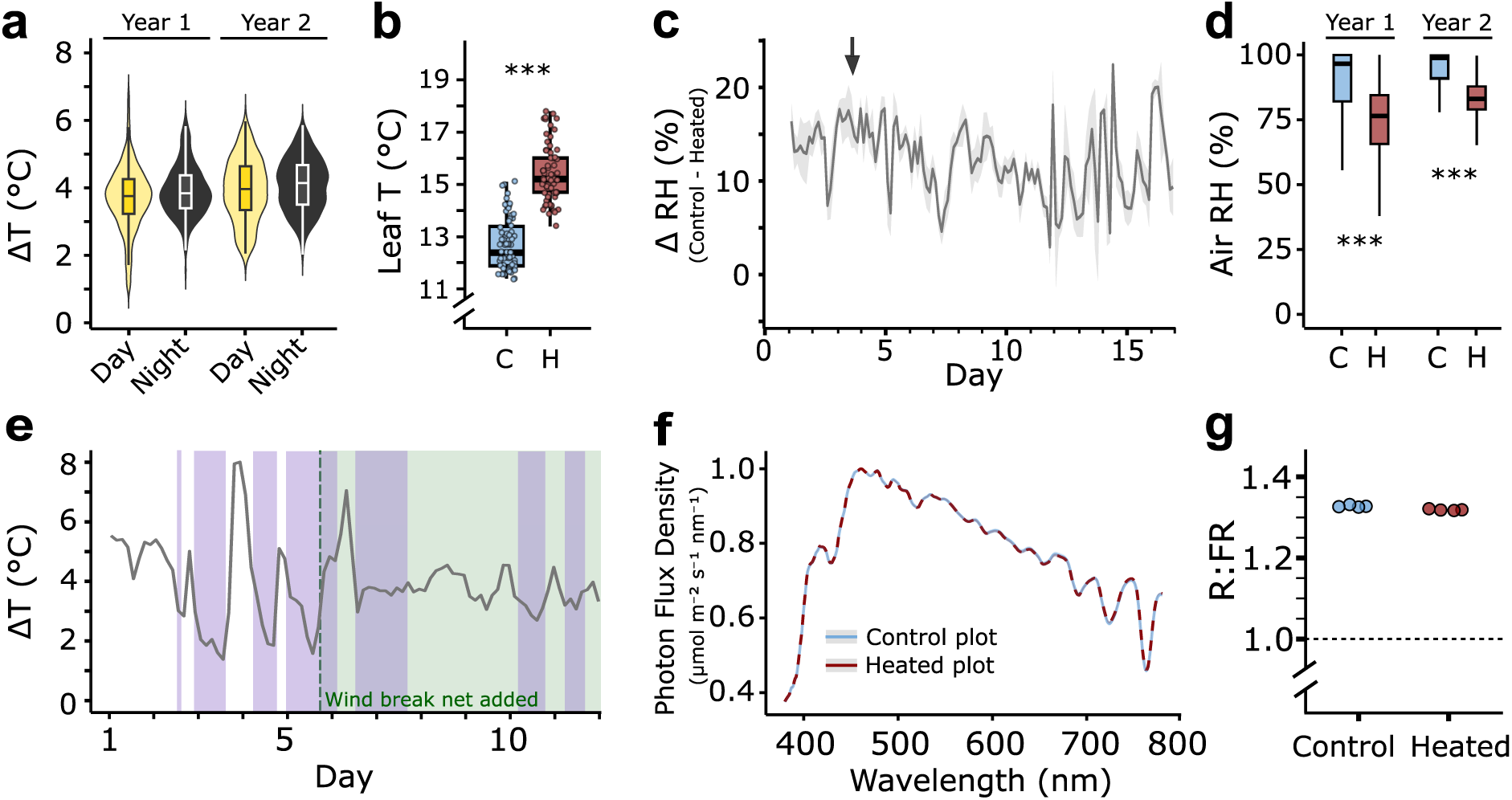
Performance of the T-FACE system. **(a)** Day ΔT versus night ΔT during Year 1 (36-day) and Year 2 (16-day) heating periods. n > 290 per group, see *Methods*. **(b)** Leaf temperature (T) measured in Year 2 (2h-single session) using MiniPAM-II. n = 69 for control, n = 70 for heated. **(c)** Differential relative humidity (Δ RH = heated – control) across a 16-day heating period in Year 2. Each datapoint represents a 3-hour time window. Arrow indicates timing of soil moisture measurement (Day 3, 29 January 2024). **(d)** Air RH in heated and control plots during Year 1 (n = 1,727 per group) and Year 2 (n = 768 per group) heating periods. **(e)** ΔT over a 12-day period in Year 1. Purple highlights indicate time intervals where wind speed was higher than 15 mph. Each datapoint represents a 3-hour window. A windbreak net was installed on the afternoon of Day 5 (4:30 pm). **(f)** Spectral profiles of heated and control plots measured using a spectrometer (n = 4 technical replicates per group). No significant difference was detected in the photosynthetically active radiation range (400–700 nm) (Functional ANOVA, *F* = 0.0012, *p* = 0.486). **(g)** red:far-red (R:FR) light ratios from the same spectrometer data (n = 4 technical replicates per group). Threshold of 1 indicates shade avoidance threshold (Vandenbussche et al., 2005; Yang & Li, 2017). **(b-d)** ***, *p* < 0.001.

To get an initial indication of heat transfer to the plants, we measured leaf temperature using the leaf surface temperature sensor of a Mini-PAM II during a single measurement session. During this session, control plot values ranged from 11.4 to 15.1 °C, while heated plot values ranged from 13.4 to 17.8 °C. On average, leaf temperature in the heated plot was 2.6 °C higher (Wilcoxon test, *W* = 185.5, *p* < 2.2 × 10⁻¹⁶) than in the control (Figure 3b). The results confirm that the system can elevate canopy temperature under the conditions tested.

To explore the effect of heating on air relative humidity (RH), we compared RH between heated and control plots across both years. Control plots had high RH, with mean values of 89.0% and 94.8% (near saturation) in Years 1 and 2, respectively, whereas heated plots showed significantly reduced RH, with means of 74.3% and 82.7% (Wilcoxon test, *W* =2,350,086 in Year 1; *W* = 513,600 in Year 2, *p*-adj < 2.2 × 10⁻¹⁶). On average, RH in heated plots was 13.4% lower than in control plots (Figure 3c-d). To determine whether the difference in RH would impact soil moisture, we assessed soil moisture in both plots using a consumer-grade 4-in-1 soil tester. This qualitative instrument is able to distinguish between dry (values between 0–2) and wet soils (values above 2). Despite differences in RH at the time of measurement (89.0 % in control plot versus 72.2% in heated plot; Figure 3c), soil moisture values were consistently within the wet range (5.7 to 8) across all areas tested, both in control and T-FACE plots (Figure S1g). This suggests that while the atmosphere became drier within the heat treatment (lower RH), the soil retained moisture.

We next assessed the effect of wind on heating efficiency with and without a windbreak net. We analysed ΔT over an 11-day period in Year 1, during which a windbreak net was installed on Day 5. Prior to installation, the mean ΔT was 4.0 °C, compared to 3.9°C after. The range of ΔT, however, decreased after windbreak installation (1.4 °C – 8.0 °C before; 2.7 °C – 7.0 °C after), particularly when wind speeds exceeded 15 mph (Figure 3e). This suggests that inclusion of a windbreak net helps maintain a more stable ΔT environment and avoids the larger fluctuations observed at wind speeds of > 15 mph.

We selected heaters that emit far-infrared radiation and which are not expected to emit PAR, which could confound experiments by altering the light environment within the heated plots. Therefore, we sought to confirm the absence of PAR by measuring photon flux density (PFD) across the 400–700 nm range using a spectrometer. The spectral profiles of heated and control plots were identical, with no significant differences in PFD (Functional ANOVA, *F* = 0.0012, *p* = 0.486), suggesting no PAR was emitted from the heaters (Figure 3f). Additionally, both treatments had red:far-red (R:FR) light ratios over 1.3, above the shade avoidance threshold of 1 (Figure 3g)(Vandenbussche et al., 2005; Yang & Li, 2017). These spectral data confirm that the infra-red heaters emitted no detectable PAR across the 400-700 nm range.

Finally, we wanted to assess whether the heat treatment caused physiological stress. We therefore quantified Fv/Fm, the maximum quantum yield of Photosystem II, as an indirect measure of overall plant health. In non-stressed plants, Fv/Fm typically reaches ∼ 0.83 (Maxwell & Johnson, 2000), although it is not uncommon to observe some variability in field-grown wheat (values between 0.80-0.83, whereas heat-stressed plants drop to 0.68-0.78 (Haque et al., 2014)). In our study, mean Fv/Fm values were 0.802 (control) and 0.814 (heated) (Wilcoxon test, *W* = 483, *p* = 0.03; Figure S1h), both within the expected range of physiological healthy (non-stressed) plants. Overall, there is no evidence of plant stress due to heating of the T-FACE plots consistent with expectations for a mild heat treatment.

### Biological responses to heat treatment

To validate the biological relevance of the T-FACE system, we measured developmental and morphological traits in wheat. We collected phenotypic data from wheat plants sown during the Year 1 experimental period (2022–2023), when the T-FACE heating system was operated from 27^th^ January to 19^th^ May, providing 97 days of effective heating (i.e., without system failure; see next section: *Technical limitations*). We selected well-characterised lines to help with interpretation of temperature responses, using two Near-Isogenic Lines (NILs) in the hexaploid spring wheat cv ‘Paragon’. The NIL pair consisted of a Paragon wildtype line carrying the wildtype photoperiod-sensitive allele of the *PHOTOPERIOD1* (*PPD1*) gene, and a *ppd1* triple null mutant which flowers later due to the absence of functional PPD1.

We phenotyped six traits to address two key questions: (i) whether the heat treatment induced phenotypic changes relative to control plots, and (ii) whether *PPD1* allelic status modulated responses to heating. The T-FACE treatment significantly decreased plant height, spike length, spikelet number, and days to heading (DTH) across both genotypes (Figure 4a-c). Within the heated plot, *PPD1* NILs headed approximately 14 days earlier, and *ppd1* NILs about 7 days earlier, compared to control conditions. Genotype also had a strong effect on heading (two-way ANOVA, *F* (1, 12) = 161, *p* < 0.0001), with *PPD1* NILs heading earlier than *ppd1* by ∼ 8 days under control and 15 days under heated conditions. This difference in magnitude under heat treatment is reflected in a significant genotype-by-treatment interaction (two-way ANOVA, *F* (1, 12) = 15, *p* = 0.0023), indicating that *PPD1* NILs are more responsive to the heat treatment than *ppd1* NILs.

**Figure 4.**
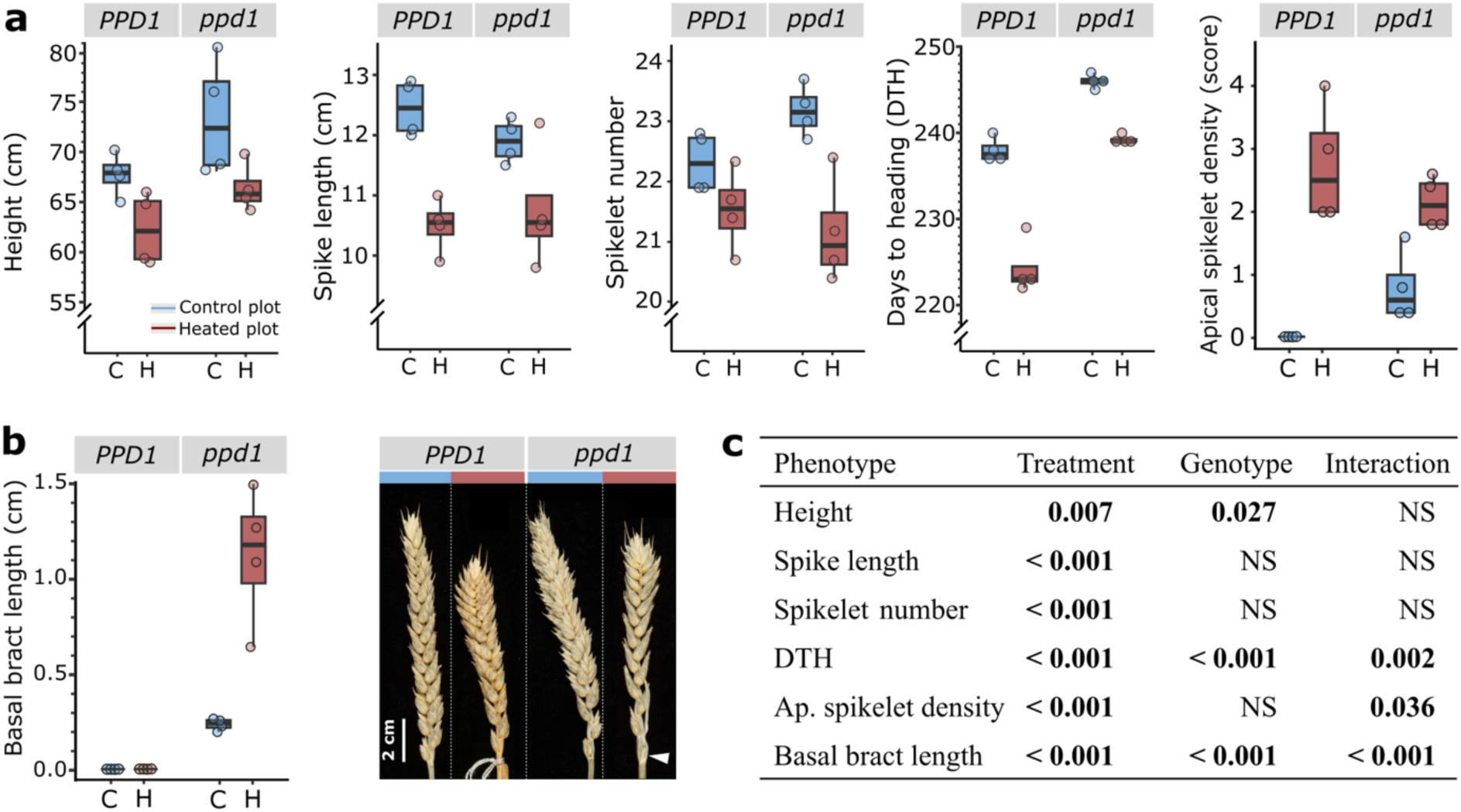
Phenotypic effects of plot warming. **(a)** Effect of Year 1 heating treatment on Paragon *PPD1* and *ppd1* NILs for crop height, spike length, spikelet number, days to heading and apical spikelet density under control (C, blue) and heated (H, red) treatments. Each datapoint represents a biological replicate. **(b)** As in (a), but for basal bract length (left) and representative spike images (right).The white arrow indicates the basal bract observed exclusively in *ppd1*. **(c)** *P*-values from two-way factorial ANOVA models assessing the effects of genotype, treatment (control vs. heated), and their interaction. Significant *p*-values < 0.05 are highlighted in bold; NS = non-significant.

We also observed additional morphological differences between treatments. Plants exposed to heating showed a significantly increased apical spikelet density (Figure 4a-b; two-way ANOVA, *F* (1, 12) = 48, *p* < 0.0001) in both genotypes. Additionally, we observed a distinct, leaf-like bract exclusively in *ppd1* NILs at the junction of the peduncle and the rachis supporting the first basal spikelet. This basal bract was absent in *PPD1* NILs irrespective of treatment and was significantly longer under heat in *ppd1* plants (two-way ANOVA, *F* (1, 12) = 24, *p* < 0.001; Figure 4b).

We observed similar trends across four additional wheat genotypes (Figure S2). All six traits had significant treatment effects (two-way ANOVA, *F* (1, 36) > 15, *p* < 0.001), with a modest reduction in plant height and spikelet number, and more significant decrease in spike length and DTH. The reduction in spike length can be partially explained by the increase in apical spikelet density, which is significantly increased under heat. This increase in apical spikelet density, however, was genotype-dependent and absent in *Triticum turanicum* under both conditions. The basal bract phenotype remained exclusive to *ppd1* NILs. Overall, the consistent phenotypic responses across genotypes indicate that the T-FACE system provides a biologically active and effective heat treatment under field conditions.

### Technical limitations of the system and costs

While the T-FACE system was effective and elicited biological changes, several technical and operational challenges were encountered. The main technical challenge encountered during the experiment was the repeated failure of the infrared heaters. In Year 1, heater failure occurred on three separate occasions, while in Year 2 it happened on five occasions (Figure S3). Each failure of an individual heater triggered an electrical earth fault, which tripped the main RCD and subsequently shut down all heaters in the hexagonal array. While the initial investment was substantial (£23,152; see Table S1), running costs are modest and primarily linked to electricity costs. During Year 1 (January to March 2023), the T-FACE system consumed an average of 124.5 kWh per day per hexagonal heater array when configured to maintain a ΔT of + 5 °C. Based on the current UK average electricity rate (as of May 2025), which ranges from £0.21 to £0.27 per kWh, the estimated daily operational cost per array is approximately £26.15. Each hexagonal heating array covers a surface area of approximately 7.77 m², resulting in an operational cost of around £3.36 per m^2^ per day. At this value, operational costs are lower than typical rates for controlled environment chambers, making the T-FACE system cost-effective for field-based temperature manipulation experiments.

## Discussion

Our system performed well overall, demonstrating that a UK-compliant free-air infrared heating array can be deployed successfully under real field conditions. Despite challenges, most notably heater reliability, we gained valuable insights into design, control, instrumentation, and costs that may support others building similar systems. Our experience highlights both the potential and the practical realities of using open-field infrared heating in temperate climates.

### Designing for system effectiveness

Consistent sensor placement was critical to the control system’s reliability, given that the heating treatment is based on average air temperature. Sensors driving the control system should be oriented consistently, ideally facing away from the sun, to ensure accurate readings and temperature regulation by the control panel. Similarly, temperature and humidity loggers should be positioned to avoid direct sun exposure on their sensor elements. While we achieved reasonably uniform heating across the hexagonal plots, we observed minor temperature gradients, likely caused by wind direction, sensor angle relative to the heaters, and localised shading by the array itself. These small spatial differences could be used creatively in future experiments, for example, to explore responses to thermal gradients.

The use of a windbreak net effectively reduced heat loss at our exposed site in Norwich U.K, though placement must be managed carefully to avoid unintended shading. Structural elements of the heater system can also cast shadows, so it is important to consider sun angle, nearby infrastructure, and the relative layout of heated and control plots. Future experiments should include a control plot with the same scaffold structure but without active heating to isolate any effects due to shading caused by the frame itself. Mirroring the number and position of temperature loggers across control and heated plots is also important to ensure accurate ΔT estimations. Stevenson screens are not suitable for ΔT calculations in infrared heating experiments, as they shield the sensors from incoming radiation. Instead, uncovered loggers should be used to assess heating efficiency, supplemented by a shielded logger for baseline ambient temperature readings.

In our experiments, we did not include a structural control plot, which could limit the interpretation of some results. While the shaded area was typically small and dynamic (Figure S4), even transient shading may introduce subtle effects upon light-sensitive traits, or interact with the warming treatment. Nonetheless, the significant differences observed across six phenotypic traits suggests that warming, rather than shading alone, was the primary driver of the measured responses.

### Considerations about heater reliability

The main technical challenge was heater reliability. After several weeks of continuous use, individual units developed faults that triggered the system’s RCD, shutting down the entire system. In retrospect, installing individual Residual Current Breaker with Overcurrent protection (RCBOs) per heater unit would have isolated these faults and prevented full system shutdown, which is something we recommend for future implementation. That said, this would not have resolved the underlying issue of the frequent failure of the heaters themselves. All seven units eventually required repair, many more than once. We found that many failures stemmed from poor internal construction: sharp stamped metal edges cutting through wire insulation, insulation surrounding the heating element was scorched or burned, and insufficient ventilation that likely led to overheating. Moisture ingress may also have contributed, and limited clearance above the heaters may have been inadequate for long-term outdoor deployment, despite an IP64 rating.

We are now working with the manufacturer to identify more robust alternatives. The Herschel team has proposed models (such as the Aspect XL3 or Advantage IR3), with potential custom modifications to improve durability and integrate with our setup. Enhancements such as protective covers, improved ventilation, and internal design adjustments may improve long-term performance. We also recommend designing heating schedules that include off-cycles, and field-testing heater models prior to full deployment. The control system itself could be further enhanced through integration with a Building Management System (BMS), enabling real-time monitoring through a graphical interface, remote adjustment of temperature setpoints, automated fault detection, and detailed data logging. While such upgrades require additional investments, they would enhance functionality, adaptability and improve user experience.

### Elevated temperature scenarios: context and breeding potential

In previous free-air warming studies, canopy-level temperature increases typically ranged from + 1 to + 3 °C utilizing 1 kW heaters (Kimball et al., 2008; Ottman et al., 2012; Zhang et al., 2020). Greater warming of + 4 °C was achieved in larger 4 m diameter plots with hexagonal arrays of high-power 8.1 kW heaters (Kimball et al., 2018). We used 2.7 kW heaters arranged in a 3-m diameter hexagonal plot. We targeted + 5 °C and measured an average warming of + 4 °C, suggesting our heaters operated near full power. This result aligns well with Kimball’s empirical expectations based on heater power and plot size (Kimball et al., 2015).

In our study, we targeted a + 5 °C warming to evaluate the upper limits of the system performance. Our observed + 4 °C warming exceeds the UKCP18 projected mean temperature rise (Jan – May) of + 2.5 °C for 2060–2080, placing our heated plot season above the 99th percentile (less than 1% likelihood). Still, such testing can be used to evaluate biological responses under extreme scenarios, including heatwaves, which are predicted to become more frequent (Perkins-Kirkpatrick & Lewis, 2020). Furthermore, deploying this warming system in UK breeding programs could allow annual screening of 5–10 elite varieties for climate resilience by exposing them to predicted (+ 2.5 °C) and severe (+ 4 °C) warming conditions. Early testing, before entry to the Recommended List, would identify critical stress responses like sterility. This could prevent detrimental scenarios like those seen for wheat cultivar Moulin in Scotland, where unforeseen environmental stress led to widespread sterility and major yield losses (Law, 1999; Hoad et al., 2018). Integrating this approach into breeding pipelines would contribute to breeding more climate-resilient crops.

## Conclusions

Despite technical challenges, this T-FACE system offers a promising, flexible and cost-effective method for studying crop responses to warming under realistic field conditions and in a UK regulation-compliant manner. The system successfully maintained targeted temperature differentials via real-time feedback control and produced measurable phenotypic effects on multiple wheat genotypes. While future iterations will benefit from improved heater durability, our experience demonstrates that field-deployable infrared warming systems are both feasible and effective for agricultural and ecological research in temperate climates. Critically, conducting these experiments locally allows researchers to capture region-specific interactions between warming, native soils, photoperiod and rainfall, factors not easily replicated by relocating studies to warmer regions. This work provides a practical, scalable foundation for simulating near-future climate scenarios in open field conditions with broad applicability across cropping systems, experimental designs, and research disciplines.

## Material and methods

### UKCP18 climate predictions data

We obtained the data from the UKCP18 Regional 12 km x 12 km projections (for Norwich coordinates: British National Grid coordinates 618000.0, 306000.0). These provide high-resolution regional climate model (RCM) outputs downscaled from global climate models (GCMs). Twelve projections were run using different plausible future climates generated through variations in physics parameters or initial conditions (model member IDs: 0, 1113, 1554, 1649, 1843, 1935, 2123, 2242, 2305, 2335, 2491, and 2868). We used the twelve models for our calculations and averaged the results of the twelve models. We selected the high-emissions *business-as-usual* scenario RCP 8.5. Temperature data downloaded included daily maximum and minimum air temperatures at 1.5 meters above ground level. For annual mean temperature change, we calculated the difference between two periods: UKCP18 2000–2020 (baseline) and UKCP18 2060–2080 (projected). Then, we focused particularly on a 112-day window (28 January to 19 May) to align with the duration of the Year 1 experiment. We compared the approximate temperature increase in our heated plot (+ 4 °C above control) with projected UKCP18 temperatures for 2060–2080. We calculated the Jan-May temperature differences relative to the 2023 control season measured with a Konnects system at the John Innes Centre field station (Church Farm), as a baseline ambient temperature reading, with daily maximum and minimum temperatures. To validate the representativeness of the 2023 reference year, we compared it to the UKCP18 2020–2040 distribution and found it to fall within the 58^th^ percentile, confirming its suitability as a current baseline year.

### Structural components of the system

We used infrared heaters (SUMMIT 2600, Herschel) to elevate canopy temperatures. To modulate heater output, we installed Sontay TT-EBB-G temperature sensors (one in the control plot and six in the heated plot). We positioned the Sontay sensors with their sensing elements facing away from direct sunlight. We commissioned EPS UK Ltd. to build the main structure, including the scaffold, heaters, and protective covers, while Pentagon Control Systems Ltd. designed and built the control panel (Supplementary Material 1).

### Thermal imaging

We captured aerial thermal images on 26 February 2024 at 14:30 h using a DJI Matrice M300 drone equipped with a DJI Zenmuse H20T thermal payload. We processed the original R-JPEG files using the DJI Thermal Analysis Tool (version 3.4.0). We manually added a qualitative temperature scale (Figure 2b).

### Air temperature and humidity measurements

We recorded air temperature (T) and air relative humidity (RH) using Tinytag Plus 2 data loggers (model TGP-4500). To survey the entire heated plot, we deployed seven loggers in Year 1 and six in Year 2. In each year, we also installed one control logger in the control plot. All Tinytag loggers were oriented with their sensing elements facing away from direct sunlight. In Year 2, we additionally installed Stevenson screen–covered Tinytag Plus 2 loggers (two in the heated plot and one in the control plot), all located at plot centre. We programmed the loggers to record T and RH data at 10- or 15-minute intervals.

We measured soil moisture at a depth of 8-10 cm using a 4-in-1 soil tester (Raintrip). To capture spatial variability, we took the measurements at eight points across the hexagonal plot. We took these measurements one day before (26 January 2024) and two days after (29 January 2024) initiating the heating treatment.

### Wind data and windbreak net

We installed a green P Dot Wolf high-density polyethylene (HDPE) knitted windbreak netting with 40% shade and approximately 50% wind reduction. We secured the net to the fence using cable ties, spaced every 1 m. The net extended 1 m vertically from the ground. We installed the net in both years. In Year 1, we installed a few days after the heating treatment started (1 February 2023; 16:30 h), while in Year 2, for the entire heating period. Exploring this period allowed us to compare heating efficiency before and after the net was added. We obtained wind speed data for Norwich from <u>timeanddate.com</u> to contextualize the observed temperature differences (Figure 3e).

### MiniPAM-II measurements: Fv/Fm and leaf temperature

We measured leaf temperature and the maximum quantum yield of Photosystem II (Fv/Fm) with a Mini-PAM II chlorophyll fluorometer (WALZ) on 19 April 2024. We began heating at 09:00 h, approximately 1 h after sunrise (only for the purpose of this measurements, not as part of the experimental heating treatment) and took measurements between 11:00 h and 14:00 h. We sampled plants randomly across the plot, ensuring homogeneous coverage of the plot. For the leaf temperature we used the leaf clip (model 2030-B) and clipped the middle section of the oldest leaf (avoiding main vein) so that the sensor was in direct contact with the abaxial (underside) surface of the leaf during each measurement. For Fv/Fm, we dark-adapted leaves using DLC-8 dark leaf clips (WALZ) for 20 minutes, then applied a saturating light pulse. We calculated Fv/Fm as (Fm - F₀)/Fm. For both leaf temperature and Fv/Fm, each reported measurement corresponds to an individual plant.

### Spectrometer measurements

We recorded photon flux density (PFD) across the 380-780 nm wavelength range and red:far-red (R:FR) ratios on 16 March 2023, between 11:00 and 11:30 h, with the heaters on. We positioned the spectrometer 0.5–1 m above the soil surface and took four measurements per treatment (control and heated plots). We took each measurement at a slightly different location across the plot.

### Experimental layout, plant material and phenotyping

We conducted the experiment at the Norwich Research Park, (NR4 7UZ; 52.626° N, 1.237° E). Each hexagonal plot included two 2.2 × 0.8 m areas and two 1.4 × 0.8 m areas (Figure S5). We arranged plants in 22 rows for the larger areas and 14 rows for the smaller areas, with nine plants per row and 10 cm spacing between plants. We hand-sowed plots with a bespoke multi-hole dibber on 21 October 2022 (Year 1) and 25 October 2023 (Year 2). Each genotype was randomized once per area, occupying 2–4 rows in each area, (Completely Randomised Design). We treated each area as one biological replicate. We left 0.3 m spacing between areas for management.

We grew six genotypes: (1) hexaploid spring wheat (*T. aestivum*) cv ‘Paragon’ wildtype (*PPD1*, *VRT-A2a* alleles), (2-3) two Paragon NILs: one carrying the *VRT-A2b* allele (Adamski et al., 2021) and one carrying a *ppd1* full knockout (Shaw et al., 2013), (4) winter wheat KWS Glasgow, (5) *Triticum durum* cultivar Cappelli, and (6) tetraploid *Triticum turanicum* accession TRI10343. Seed was sourced from the Germplasm Resource Unit (GRU), John Innes Centre.

We applied fertiliser and agrochemical treatments at recommended rates throughout the trial period to support crop growth and protect against pests and diseases (see *Supporting Data* for details).

We phenotyped six traits: days to heading (days from sowing until at least 50% of plants had half of the spikes emerged from the flag leaf sheath, GS55), plant height (from soil surface to the tip of the spike, excluding awns; mean of a 5-plant subsample), spike length (from the basal spikelet to the tip of the spike, excluding awns; mean of a 5-ear subsample), spikelet number (including both fertile and infertile spikelets; mean of a 5-10 ear subsample), apical spikelet density (using a 0–4 scoring scale, where 0 indicated sparse spikelets and 4 indicated a very dense distribution in the upper 50% of the spike; mean of a 5-ear subsample), and basal bract length (from the base to the tip of the extra bract; mean of a 5-10 ear subsample). Due to high pathogen pressure (aphids and mildew) during the second year, we did not include phenotypic measurements for Year 2.

### Data processing and statistical analysis

We conducted all analyses in *R* (version 4.3.3), using *tidyverse* (*dplyr*, *tidyr*, *ggplot2*, and *readr*) for data wrangling and plotting; *lubridate* for handling date-time formats; *MBA* for spatial interpolation; base R, *broom* and *rstatix* for statistical testing and its tidying; *viridis* for colour scales; *grid* and *gridExtra* for arranging graphical outputs; and *purrr* for functional programming. We prepared tables using *knitr*. All the scripts and data are available (https://github.com/IsabelFaci/r-TFACE-manuscript-2025.git).

To explore ΔT and ΔRH data, we exported 10- or 15-minute interval data using Tinytag Explorer software and averaged it into 30-minute intervals. For ΔT and ΔRH timeline and distribution plots (Figures 2a, 2c-d, 3a, 3c, 3d, S3), we calculated ΔT or ΔRH at each 30-min interval as the difference between the average temperature of the heated plot loggers (L1–L7 in Year 1; L1–L6 in Year 2) and a single control plot logger. We selected the period shown in each figure based on its specific purpose: the entire heating period for both years, including heater failures (Figure S3), a 36-day (1 March 2023 to 5 April 2023) continuous heating period in Year 1 (Figures 2c, 3a, 3d), 16-day (27 January 2024 to 11 February 2024) continuous heating period in Year 2 (Figures 2a, 2d, 3a, 3c-d). For the distribution plots (Figures 2c–d and 3a), we excluded negative ΔT values, which we assumed reflected logger errors (0.81% of the total data for Figure 2c, 0.02% for 2d, and 0.44% for 3a). We then trimmed each logger’s data to the 0.5 to 99.5^th^ percentile range to remove outliers. The final sample sizes per group were: Figure 2c (n = 1727), 2d (n = 768), and 3a (n > 200). Together, these filters captured over 98% of the original data for all three figures. For ΔT contour plots (Figure 2c-d), we averaged each logger’s ΔT for the same period as the respective distribution plot. We represented this average at its approximate field coordinates. Because areas outside the heated hexagon experience no warming, we added four corner points outside the hexagon with ΔT = 0. To visualize spatial variation, we applied multilevel B-spline approximation (MBA) using the R package *MBA*, well-suited for interpolating smooth surfaces from a limited number of irregularly spaced points. Finally, we clipped the figure to the hexagonal plot boundary so that only the heated-plot area is shown (corner points remain in code but are not visible in the published main figure).

To compare loggers covered and uncovered with Stevenson screens (Figure S1f), we calculated ΔT using only centrally located loggers: for uncovered comparisons, we used heated logger L1 and one control logger; for covered comparisons, we used the average of two central heated loggers and one control logger. From MiniPAM-II measurements, we included 99% of the data for leaf temperature (Figure 3b) and Fv/Fm (Figure S1h), removing outliers within treatment groups using percentile-based trimming. To analyse spectral photon flux density (PFD) curves (Figure 3f), we used functional data analysis (FDA) in the *fda.usc* package. We treated each spectral curve as a functional observation and applied a one-way functional ANOVA (*fanova.onefactor()*) to compare groups, using 1000 permutation resamples to test for treatment differences. To assess the effects of heat treatment and genotype on phenotypic traits (Figures 4, S2), we conducted two-way ANOVAs using the base R function *aov()* for each trait separately. The model included genotype, treatment, and their interaction as fixed factors.

For statistical comparisons (Figures 3a, 3b, 3d, S1f, S1h), we first assessed normality using the Shapiro–Wilk test (*shapiro.test()*) and then used Wilcoxon Rank-Sum tests (*wilcox.test()*) when the data did not meet normality assumptions. We applied the Benjamini–Hochberg correction (*p.adjust(method = “BH”)*) for multiple testing (Figure 3a).

## Supporting information

Supporting Data Year 1

Supporting Data Year 2

## Acknowledgements

We thank Dr. Bruce Kimball for his advice on the T-FACE setup at the start of the project; Prof. Stephen Dorling for his guidance and expertise on working with UKCP18 data; and Dr. Martin Vickers and Phil Robinson for their assistance with drone-based thermal imaging. We thank the JIC Field Experimentation team, Tobin Florio, Sophie Eade, and Pamela Crane for technical support in field experiments. We are grateful to Dr Noam Chayut (JIC GRU) for providing germplasm. This work was supported by the Biotechnology and Biological Sciences Research Council (BBSRC) through the Delivering Sustainable Wheat (BB/X011003/1) and Building Robustness in Crops (BB/X01102X/1) Institute Strategic Programmes; and the European Research Council (ERC-2019-COG-866328).

## Data and code availability

All data and codes are available at: https://github.com/IsabelFaci/r-TFACE-manuscript-2025.git

## Author information

**IF**: conceptualization, methodology, data curation, software, formal analysis, investigation, visualization, project administration, supervision, and writing—original draft; **PR**: conceptualization, methodology, resources, supervision and investigation; **JS**: methodology, investigation and resources; **MH**: methodology, investigation, validation and resources; **DP**: methodology, investigation and resources; **AD**: methodology, supervision and resources; **CU**: conceptualization, funding acquisition, methodology, project administration, supervision, validation and writing—review and editing.

## Competing interests

The authors declare no competing interests.

## Supplementary figure legends

**Figure S1. (a)** Aluminium cover protecting the heater (left) and its design schematic (right). **(b)** Adjustable handrail brackets connecting the aluminium pole to the heater cover. **(c)** Internal view of the control panel. **(d)** Warning signage indicating *Hot surface - Do not touch* (left) and thermal image showing heater and cover temperatures (right). **(e)** Locked perimeter enclosing T-FACE system and control plots. **(f)** Comparison between uncovered loggers and Stevenson-screen covered loggers in recorded temperature differences. ΔT was calculated as the difference between heated and control plots within each cover type: ΔT covered (white) = T heated-covered – T control-covered; ΔT uncovered (yellow) = T heated-uncovered – T control-uncovered. The loggers in the heated plot used for this calculation were positioned at the centre, 10 cm apart, as shown in the right-hand image. *n* = 768 per group. Wilcoxon rank-sum test, *W* = 30,865, *p* < 2.2 × 10⁻¹⁶**. (g)** Soil moisture in Year 2 control and heated plots one day before and two days after the heating treatment started. Hexagonal graphs represent: (top) control and heated plot (outlined by six rectangles representing heaters) before the heating treatment; (bottom) control and heated plots during the heating treatment. The scale shows the qualitative moisture range used by the 4-in-1 soil tester (dry = 0–2; wet > 2). **(h)** Comparison of the maximum quantum yield of Photosystem II (Fv/Fm) between control (*n* = 26) and heated (*n* = 53) plots, measured using the Mini-PAM II fluorometer. Measurements were taken after dark-adapting leaves for 20 minutes using a leaf clip, during a single 2-hour session. The plot displays 99% of the data per group. Wilcoxon test, *W* = 482.5, *p* = 0.0316.

**Figure S2.** Phenotypic effects of Year 1 heating treatment (as in Figure 4) for six genotypes used in this study. Each datapoint is a biological replicate. Table shows *p*-values from two-way factorial ANOVA models assessing the effects of genotype, treatment (control vs. heated). Significant *p*-values (< 0.05) are highlighted in bold.

**Figure S3.** Air temperature and ΔT over time across the entire heating periods (including times when the system tripped due to heater failure). These periods are marked with white squares in the top panels: **(a)** three off-periods in Year 1, and **(b)** five off-periods in Year 2. Day corresponds to days from start of temperature treatment.

**Figure S4.** Images illustrating the variation in dynamic shading across the plots throughout the year and at different times of day.

**Figure S5.** Experimental layout for Year 1, following a completely randomised design (CRD), for both control and heated plots. Lines within each plot indicate rows, and each dot within a row represents an individual plant (nine per row).

**Table S1.** Experimental costs for the HSE-compliant T-FACE system with real-time control designed in this study. Costs include main components prices (scaffold, control panel, heater covers, and heaters) as well as additional equipment such as temperature and relative humidity (RH) loggers.

## Notes

https://github.com/IsabelFaci/r-TFACE-manuscript-2025.git

### Competing Interest Statement

The authors have declared no competing interest.

