## Supplementary Figures and Tables for "Design and deployment of a regulation-compliant infrared heating system for UK field trials"

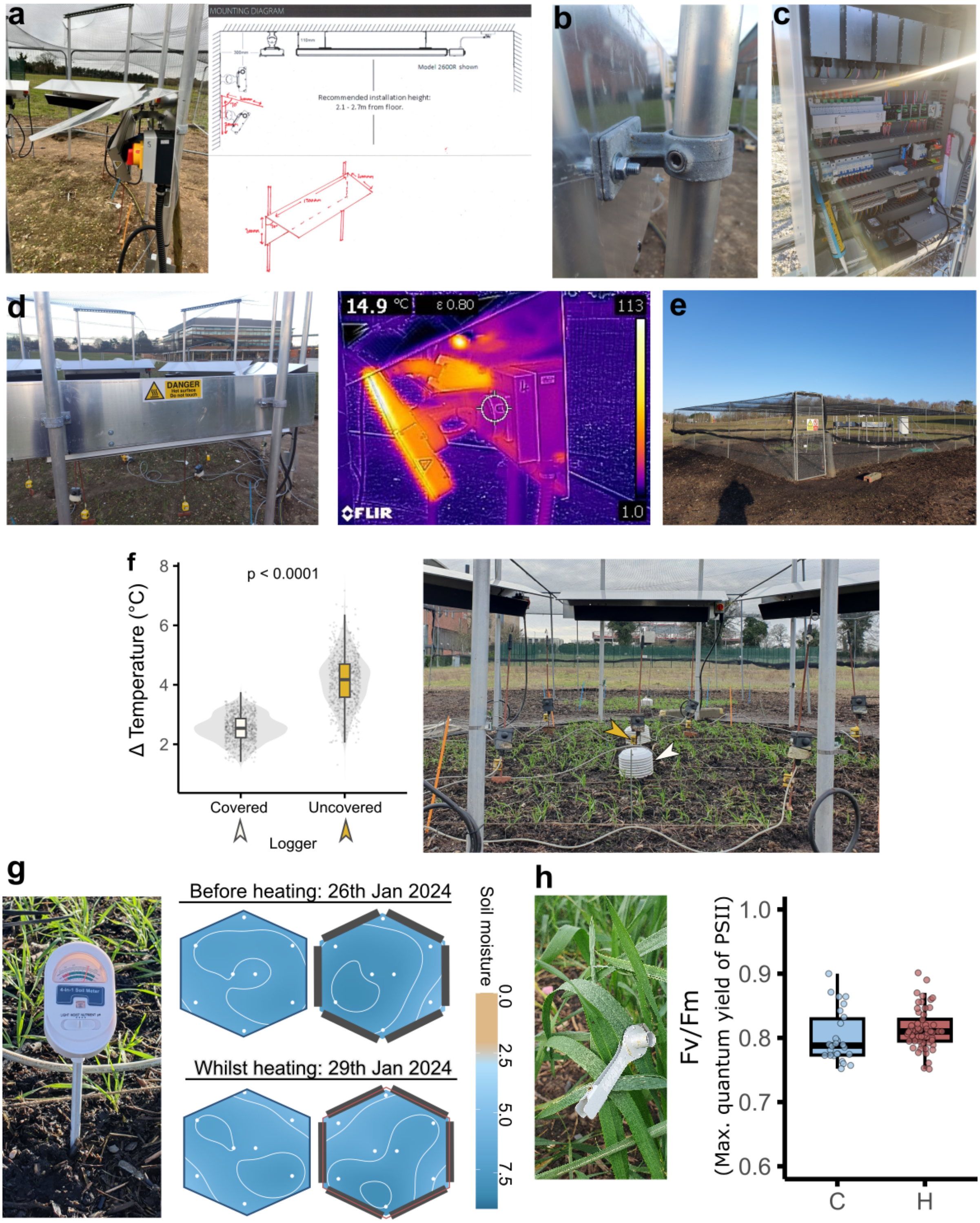
a
MOUNTING DIAGRAM
Model 2600R shown
Recommended installation height:
2.1 - 2.7m from floor.
b
c
d
DANGER
14.9 °C
ε 0.80
113
1.0
FLIR
e
f
p < 0.0001
Δ Temperature (°C)
Covered
Uncovered
Logger
g
Before heating: 26th Jan 2024
Whilst heating: 29th Jan 2024
Soil moisture
0.0
2.5
5.0
7.5
h
Fv/Fm
(Max. quantum yield of PSII)
1.0
0.9
0.8
0.7
0.6
C
H

**Figure S1.**

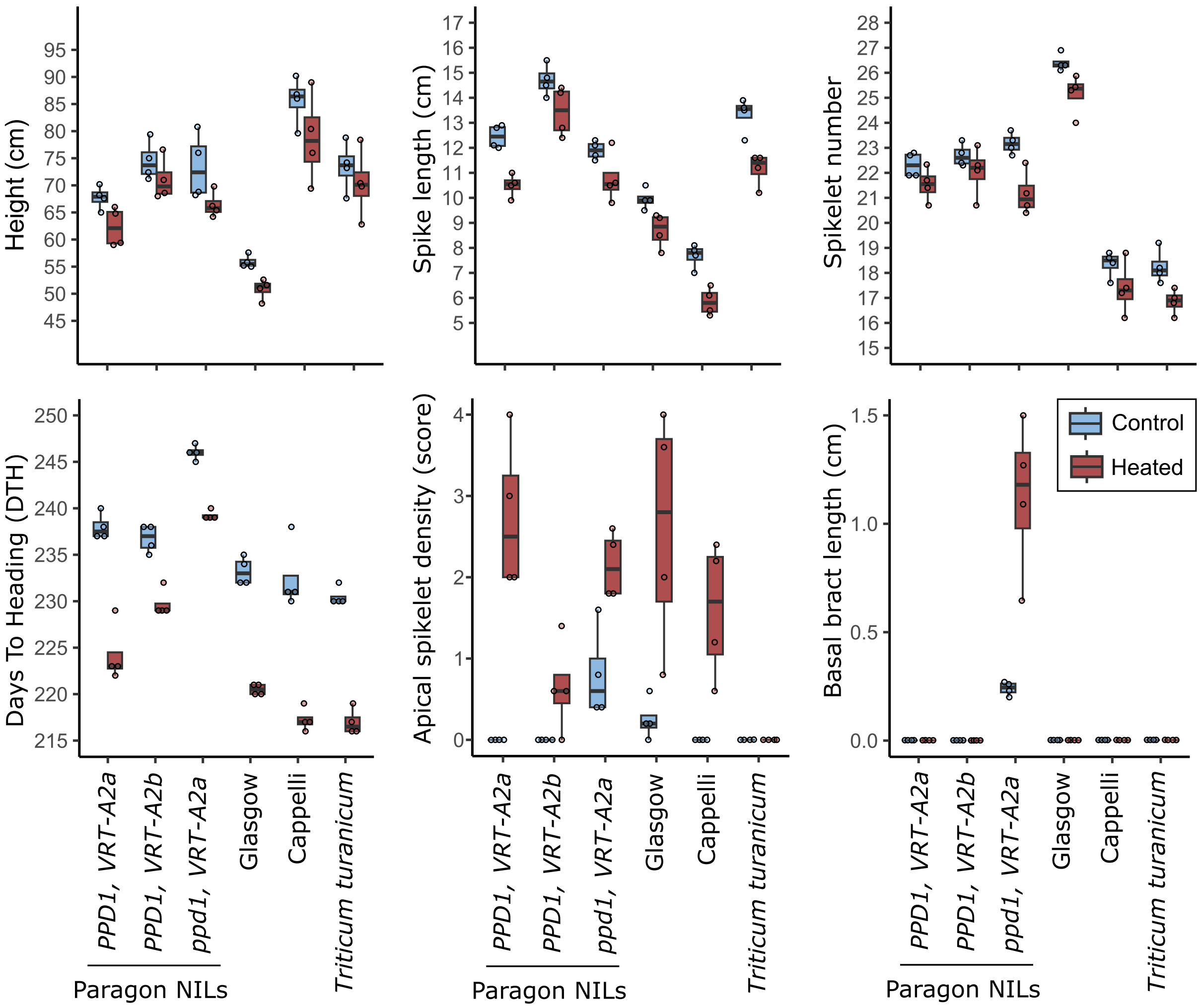


| Phenotype | Treatment | Genotype | Interaction |
|---|---|---|---|
| Height | < **0.001** | < **0.001** | NS |
| Spike length | < **0.001** | < **0.001** | NS |
| Spikelet number | < **0.001** | < **0.001** | NS |
| DTH | < **0.001** | < **0.001** | < **0.001** |
| Apical spikelet density | < **0.001** | < **0.001** | < **0.001** |
| Basal bract length | < **0.001** | < **0.001** | < **0.001** |

**Figure S2.**

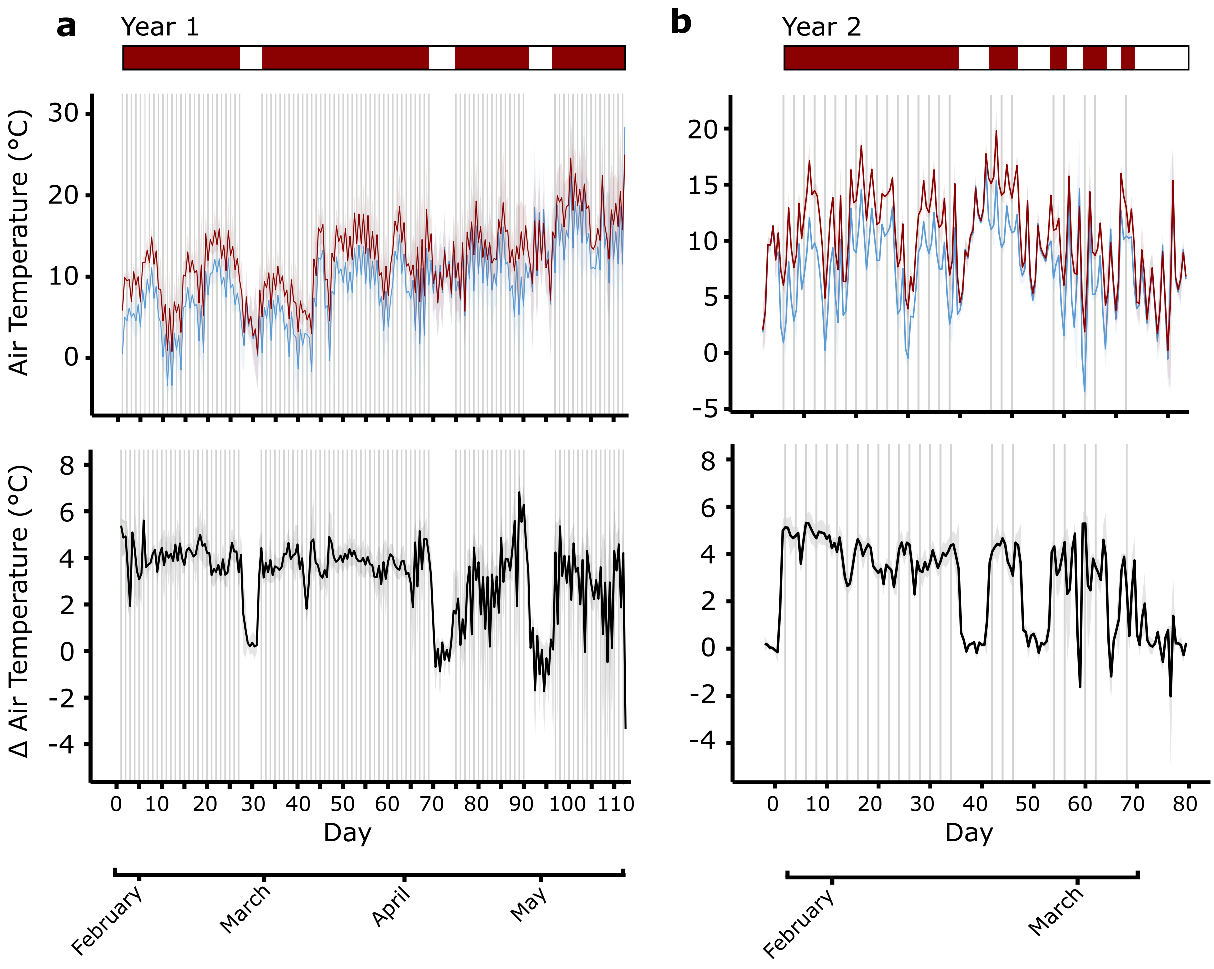
a
Year 1
Air Temperature (°C)
Δ Air Temperature (°C)
Day
February
March
April
May
b
Year 2
Day
February
March

**Figure S3.**

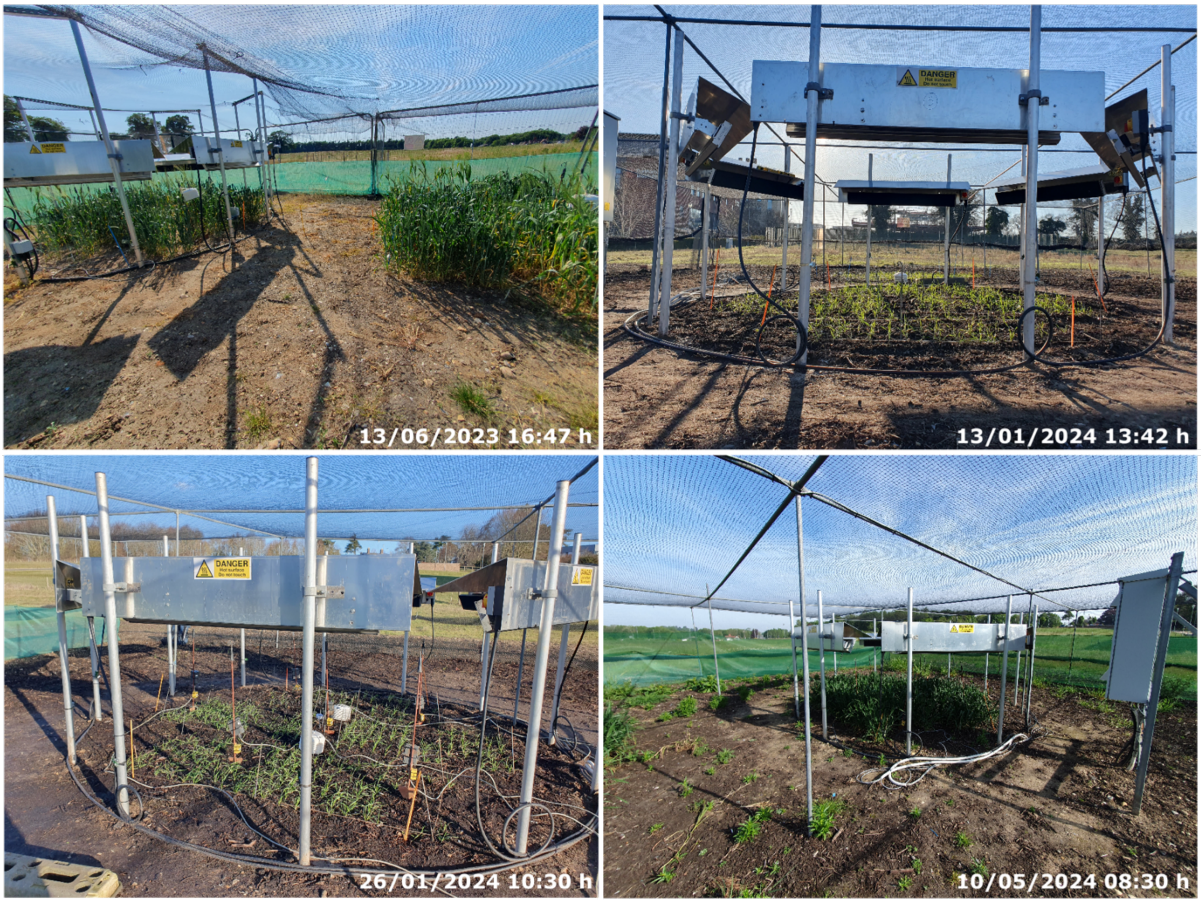


**Figure S4.**

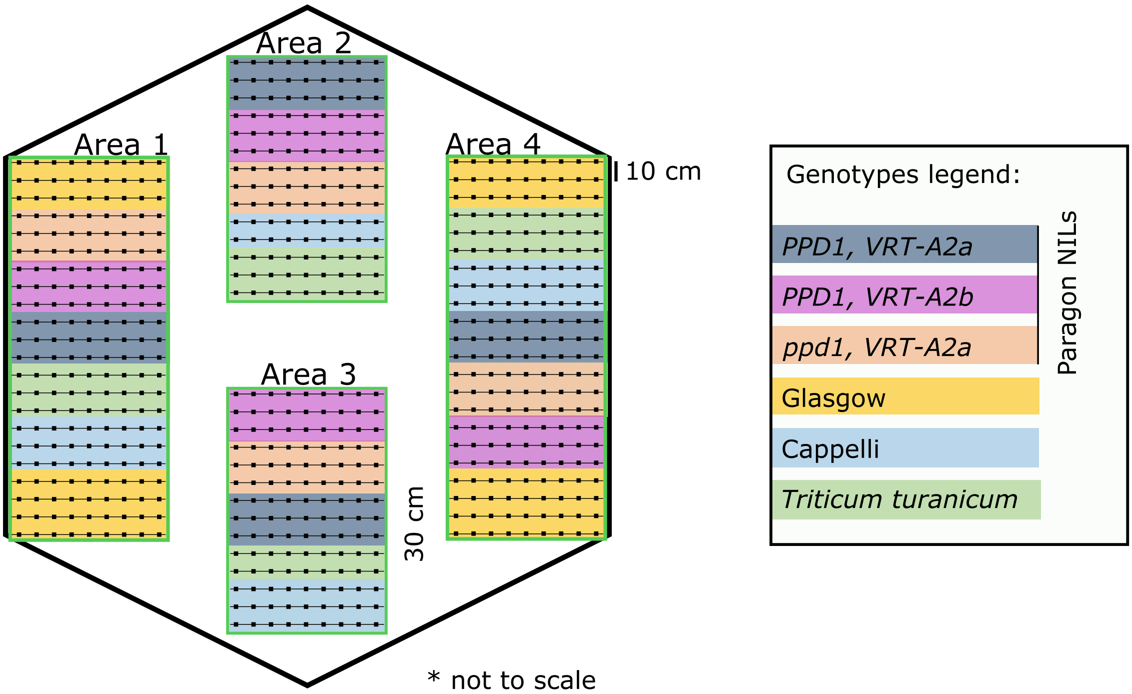


**Figure S5.**

**Table S1.**

| Component | Provider | Service/components provided | Unit price | Unit quantity | Total excl. VAT | Total incl. VAT |
|---|---|---|---|---|---|---|
| **Scaffold** | EPS | Provide scaffold and set up scaffold, heaters and covers | NA | NA | £7,045.00 | £8,454.00 |
| **Control panel** | Pentagon | Provide and set up control panel | NA | NA | £7,613.00 | £9,135.60 |
| **Covers** | bartech-services | Provide aluminium heaters covers | NA | NA | £740.62 | £888.74 |
| **Heaters** | REXEL Ltd | IR-heaters Herschel Summit 2,600W | £329.25 | 7 | £2,304.75 | £2,765.70 |
| **T and RH loggers** | Gemini Data Loggers Ltd | Tinytags plus 2 TGP-4500 | £159.00 | 10 | £1,590.00 | £1,908.00 |
| | | | | | **Total incl. VAT** | **£23,152.04** |
