## Supplementary Material 1 for "Design and deployment of a regulation-compliant infrared heating system for UK field trials"

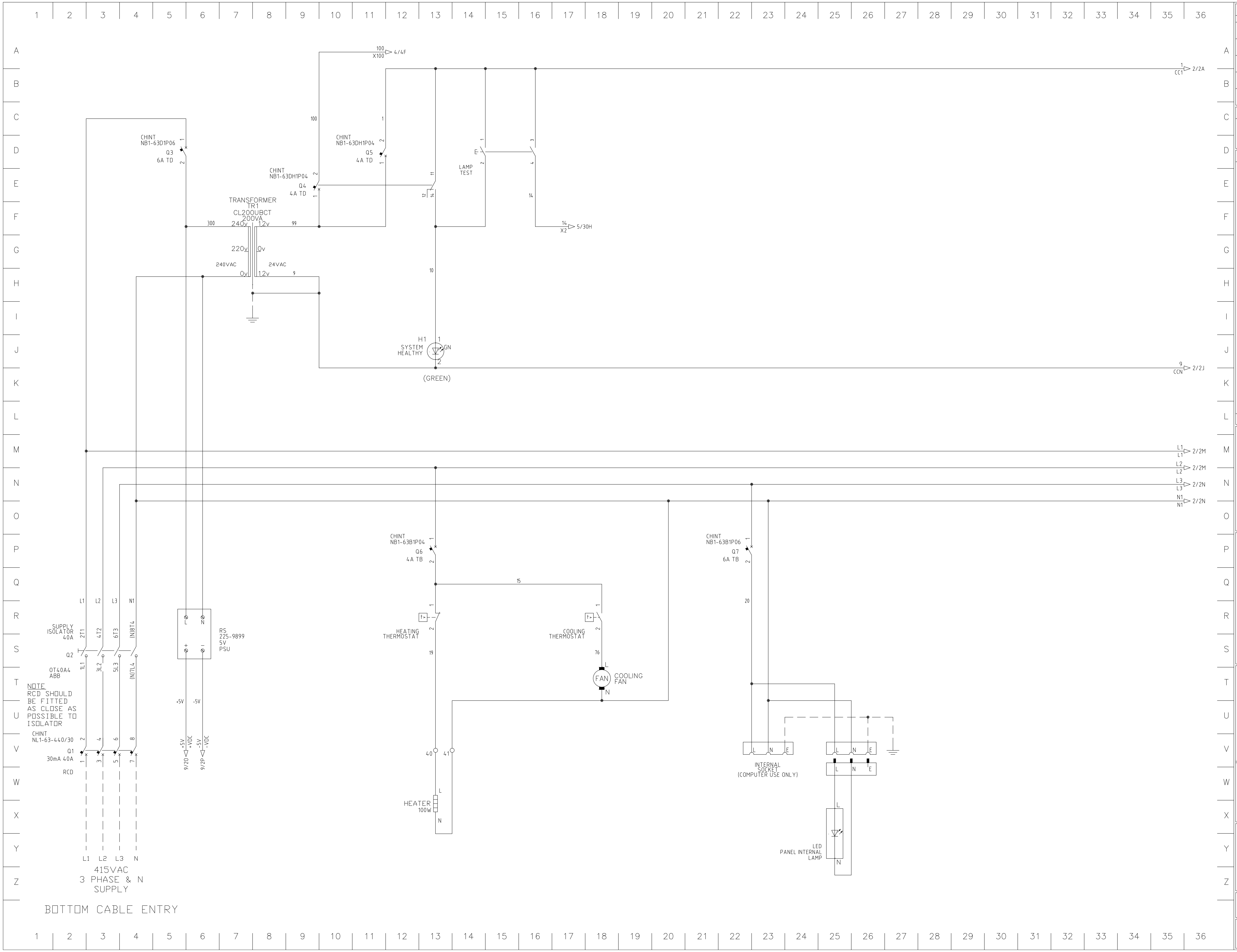

### REVISION HISTORY

| REV | DESCRIPTION | ENG | DATE |
|---|---|---|---|
| A | Client approval | GJC | 09/11/2022 |
| M | Manufacture | GWW | 28/11/2022 |
| T | Test | GWW | 05/12/2022 |
| T1 | Heater switching changed | GJC | 03/02/2023 |

### REVISION LEGEND

| | |
|---|---|
| A | MODIFICATIONS DURING CLIENT APPROVAL |
| M | MODIFICATIONS DURING MANUFACTURE |
| T | MODIFICATIONS DURING TESTING |
| AF | MODIFICATIONS DURING COMMISSIONING |

DRAWING INFORMATION

CAD Ref: 1

Pentagon
Control Systems
UNIT 8
BURNET ROAD
NORWICH
NR3 2BS

HEATING & VENTILATION CONTROL SYSTEMS

CLIENT
N.B.I.P Ltd
John Innes Centre
Norwich Research
Park, Colney
Norwich NR4 7UH

PROJECT DESCRIPTION
John Innes Centre
Field Heating Trials

DRAWING DESCRIPTION
Control Panel
Wiring Diagram

DRAWING SHEET NUMBER
1 OF 5

PROJECT NUMBER
P32700_CP001

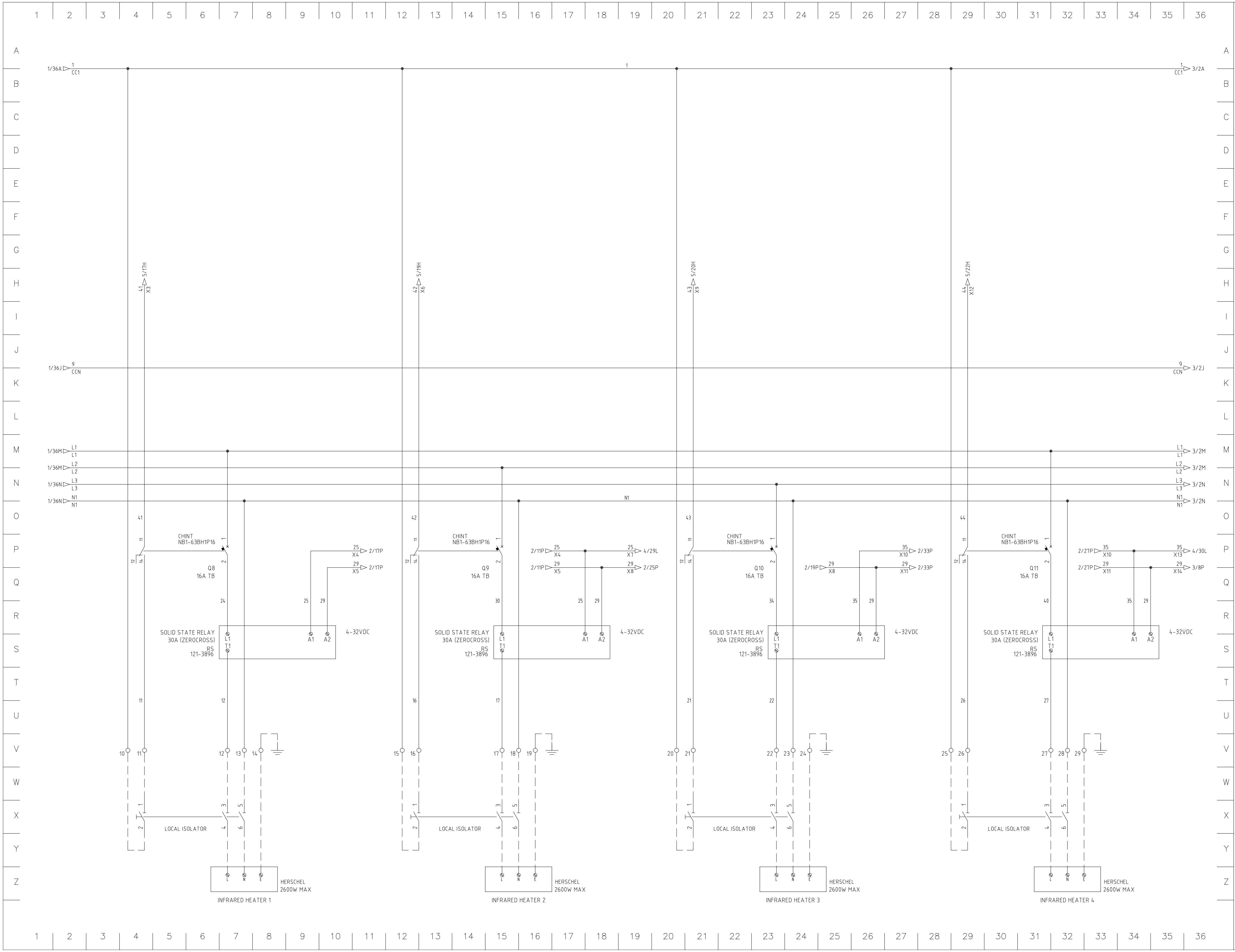

| REVISION HISTORY | | | |
|---|---|---|---|
| REV | DESCRIPTION | ENG | DATE |
| A | Client approval | GJC | 09/11/2022 |
| M | Manufacture | GWW | 28/11/2022 |
| T | Test | GWW | 05/12/2022 |
| T1 | Heater switching changed | GJC | 03/02/2023 |

| REVISION LEGEND | |
|---|---|
| A | MODIFICATIONS DURING CLIENT APPROVAL |
| M | MODIFICATIONS DURING MANUFACTURE |
| T | MODIFICATIONS DURING TESTING |
| AF | MODIFICATIONS DURING COMMISSIONING |

DRAWING INFORMATION

CAD Ref: 2

Pentagon Control Systems
UNIT 8
BURNET ROAD
NORWICH
NR3 2BS

HEATING & VENTILATION CONTROL SYSTEMS

CLIENT
N.B.I.P Ltd
John Innes Centre
Norwich Research
Park, Colney
Norwich NR4 7UH

PROJECT DESCRIPTION
John Innes Centre

Field Heating Trials

DRAWING DESCRIPTION

Control Panel
Wiring Diagram

DRAWING SHEET NUMBER
2 OF 5

PROJECT NUMBER
P32700_CP001

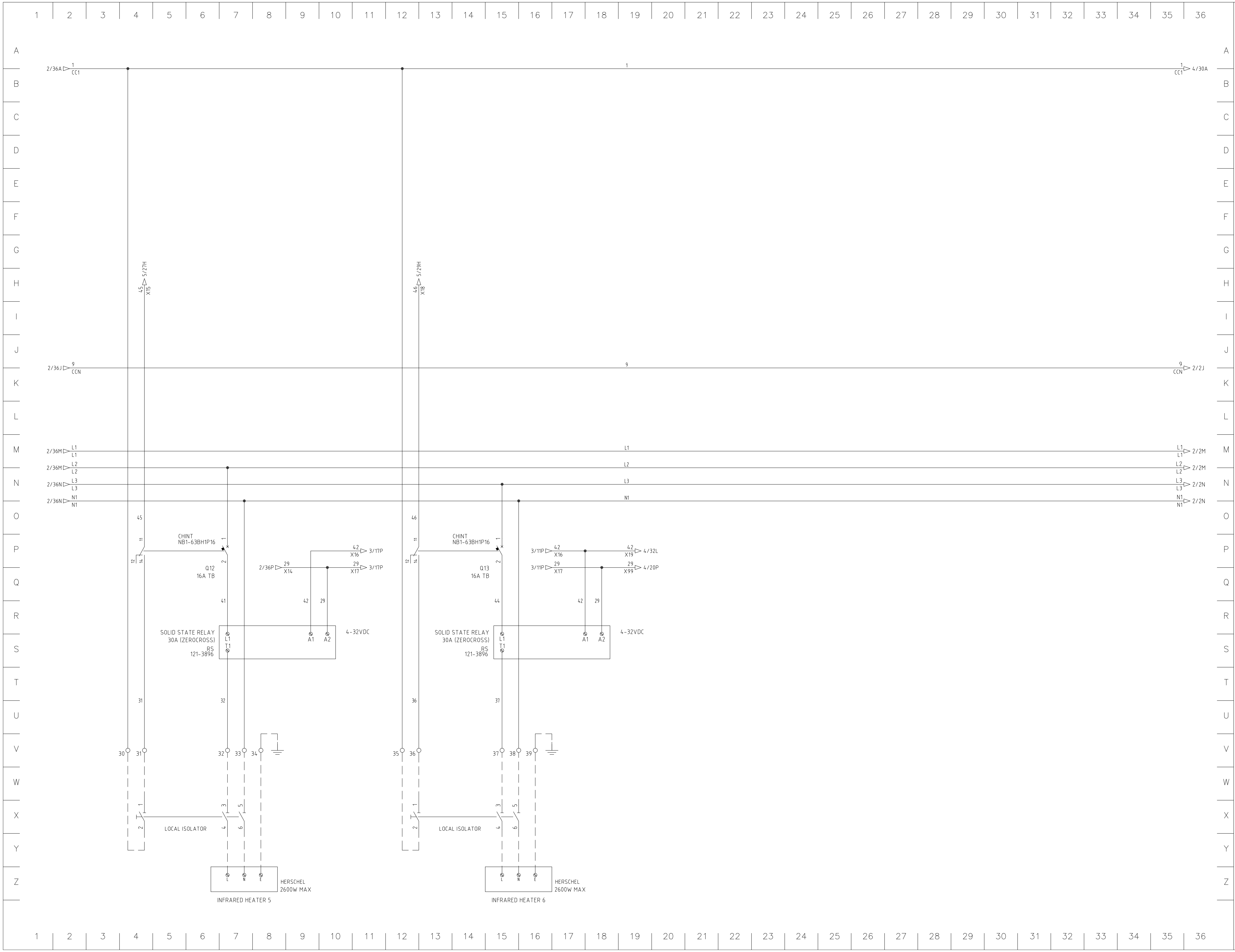

### REVISION HISTORY

| REV | DESCRIPTION | ENG | DATE |
|---|---|---|---|
| A | Client approval | GJC | 09/11/2022 |
| M | Manufacture | GWW | 28/11/2022 |
| T | Test | GWW | 05/12/2022 |
| T1 | Heater switching changed | GJC | 03/02/2023 |

### REVISION LEGEND

A MODIFICATIONS DURING CLIENT APPROVAL
M MODIFICATIONS DURING MANUFACTURE
T MODIFICATIONS DURING TESTING
AF MODIFICATIONS DURING COMMISSIONING

### DRAWING INFORMATION

CAD Ref: 3

Pentagon
Control Systems
UNIT 8
BURNET ROAD
NORWICH
NR3 2BS

HEATING & VENTILATION CONTROL SYSTEMS

CLIENT
N.B.I.P Ltd
John Innes Centre
Norwich Research
Park, Colney
Norwich NR4 7UH

PROJECT DESCRIPTION
John Innes Centre
Field Heating Trials

DRAWING DESCRIPTION
Control Panel
Wiring Diagram

DRAWING SHEET NUMBER
3 OF 5

PROJECT NUMBER
P32700_CP001

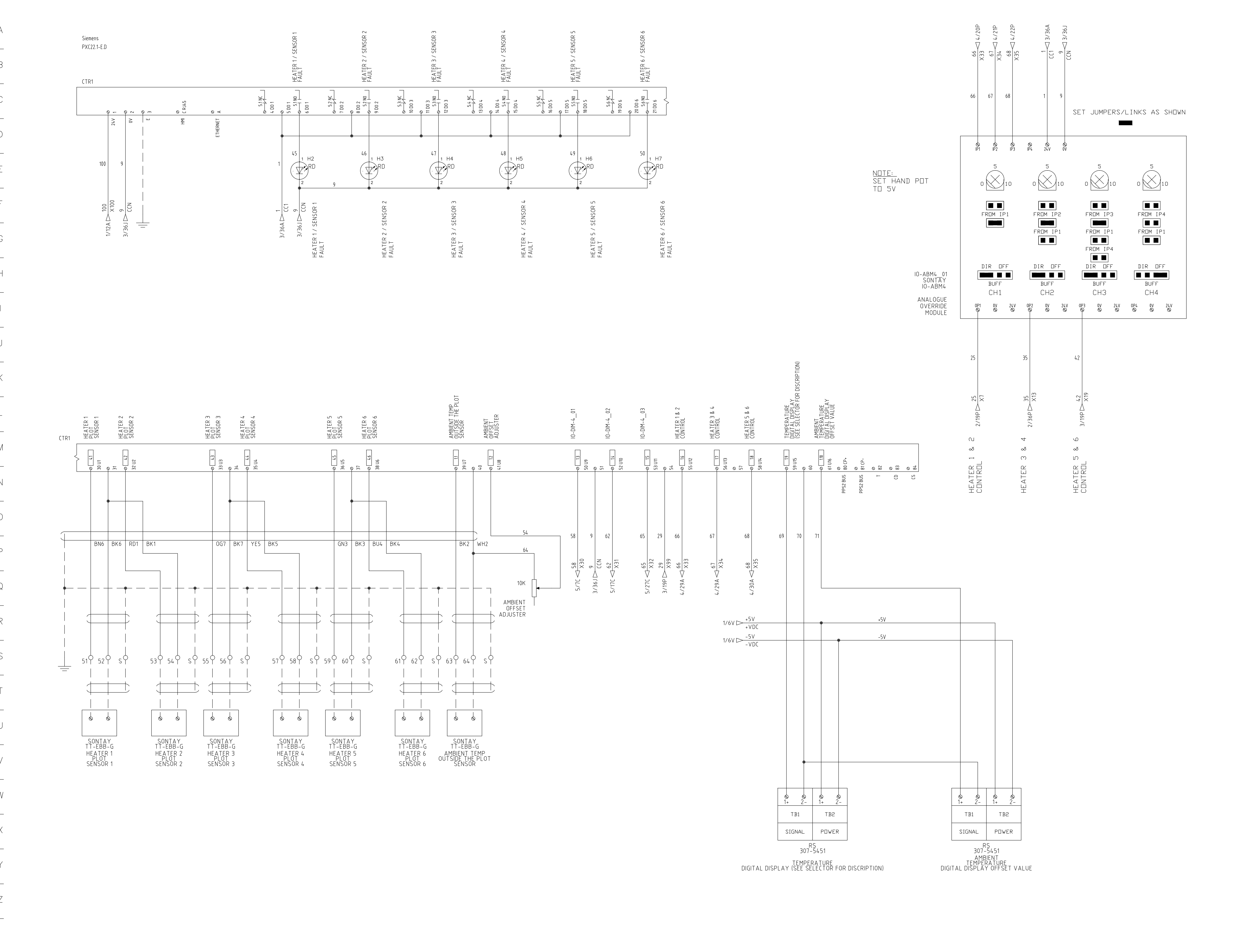

### REVISION HISTORY

| REV | DESCRIPTION | ENG | DATE |
|---|---|---|---|
| A | Client approval | GJC | 09/11/2022 |
| M | Manufacture | GWW | 28/11/2022 |
| T | Test | GWW | 05/12/2022 |
| T1 | Heater switching changed | GJC | 03/02/2023 |

### REVISION LEGEND

| | |
|---|---|
| A | MODIFICATIONS DURING CLIENT APPROVAL |
| M | MODIFICATIONS DURING MANUFACTURE |
| T | MODIFICATIONS DURING TESTING |
| AF | MODIFICATIONS DURING COMMISSIONING |

DRAWING INFORMATION

CAD Ref: 4

Pentagon Control Systems
UNIT 8
BURNET ROAD
NORWICH
NR3 2BS

PCS
HEATING & VENTILATION CONTROL SYSTEMS

CLIENT
N.B.I.P Ltd
John Innes Centre
Norwich Research
Park, Colney
Norwich NR4 7UH

PROJECT DESCRIPTION
John Innes Centre
Field Heating Trials

DRAWING DESCRIPTION
Control Panel
Wiring Diagram

DRAWING SHEET NUMBER
4 OF 5

PROJECT NUMBER
P32700_CP001

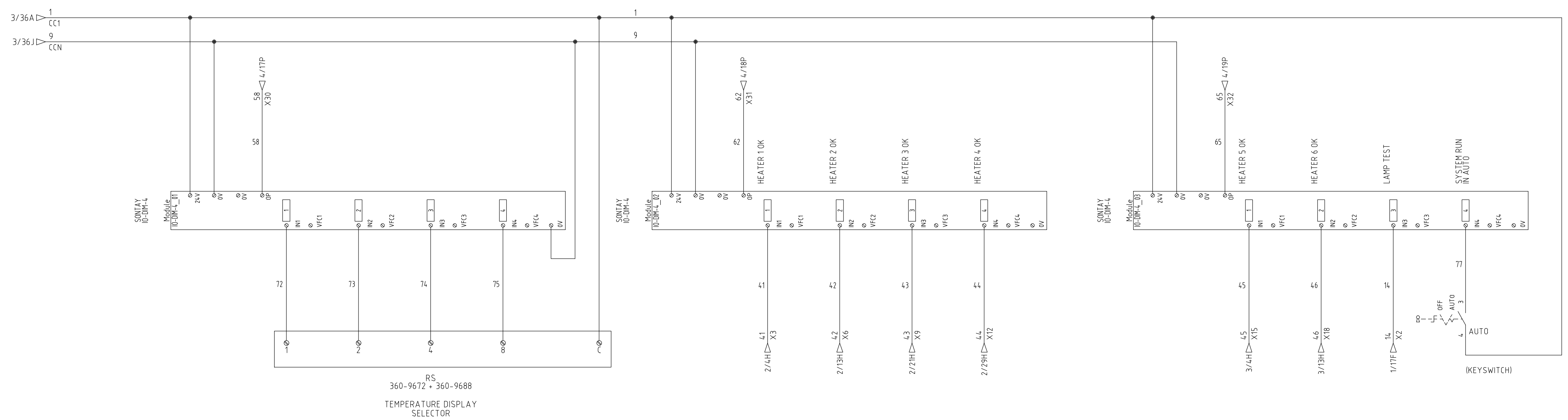

0-9 BCD PUSHWHEEL SWITCH

0-AMBIENT TEMPERATURE
1-HEATER 1 SENSOR
2-HEATER 2 SENSOR
3-HEATER 3 SENSOR
4-HEATER 4 SENSOR
5-HEATER 5 SENSOR
6-HEATER 6 SENSOR
7-MINIMUM PLOT SENSOR TEMPERATURE
8-MAXIMUM PLOT SENSOR TEMPERATURE
9-AVERAGE PLOT SENSOR TEMPERATURE

REVISION HISTORY

| REV | DESCRIPTION | ENG | DATE |
|---|---|---|---|
| A | Client approval | GJC | 09/11/2022 |
| M | Manufacture | GWW | 28/11/2022 |
| T | Test | GWW | 05/12/2022 |
| T1 | Heater switching changed | GJC | 03/02/2023 |

REVISION LEGEND

| | |
|---|---|
| A | MODIFICATIONS DURING CLIENT APPROVAL |
| M | MODIFICATIONS DURING MANUFACTURE |
| T | MODIFICATIONS DURING TESTING |
| AF | MODIFICATIONS DURING COMMISSIONING |

DRAWING INFORMATION

CAD Ref: 5

Pentagon
Control Systems
UNIT 8
BURNET ROAD
NORWICH
NR3 2BS

HEATING & VENTILATION CONTROL SYSTEMS

CLIENT
N.B.I.P Ltd
John Innes Centre
Norwich Research
Park, Colney
Norwich NR4 7UH

PROJECT DESCRIPTION
John Innes Centre
Field Heating Trials

DRAWING DESCRIPTION
Control Panel
Wiring Diagram

DRAWING SHEET NUMBER
5 OF 5

PROJECT NUMBER
P32700_CP001
