## Supplementary Material 2 for "Design and deployment of a regulation-compliant infrared heating system for UK field trials"

| | |
|---|---|
| **JOHN INNES CENTRE<br>STANDARD OPERATING PROCEDURE** | |
| **TITLE:** Adjusting the height of heaters in the free-air heating arrays (T-FACE system: Temperature Free-Air Controlled Enhancement) | |
| **APPLIES TO STAFF IN:** Cristobal Uauy's team | |
| **HEALTH & SAFETY INFORMATION INCLUDED:** **YES** / NO | |
| **REFERENCE No:** cg_cu_005 | **VERSION No:** 001 |
| **DATE EFFECTIVE:** 16.02.2023 | **REVIEW DATE:** 15.02.2025 |
| **AUTHOR:** Isabel Faci-Gomez | **APPROVED BY:** Cristobal Uauy |
| **H&S and QA AUTHORISATION:** Laurence Tomlinson | **DATE ADDED TO QA DATABASE:** 16.02.2023 |

This is a controlled document maintained on electronic media. When appearing in paper form it should be checked against the master on the QA database to ensure that the current version is being used. This copy was printed on 16 February 2023.

### 1 PURPOSE OF PROCEDURE/METHOD AND ITS SCOPE

This SOP details the correct procedures for adjusting the height of heaters in the free-air heating arrays (T-FACE system: Temperature Free-Air Controlled Enhancement). Authorized personnel must receive training in the procedure. All personnel must read this SOP as part of the training. Please fill in details of this and any problems that have been encountered with the run and if necessary, report this to the H&S team.

### 2 EQUIPMENT AND REAGENTS NEEDED

**Equipment:**

- Hexagon Allen key
- Nitrile foam gloves

### 3 STEPS IN PROCEDURE

1) Turn off the T-FACE system in the control panel (key needed). Let the heaters cool down for ~60 minutes. Note: The heater bodies are very hot (>180 °C) when the system is ON, but this heat does not conduct to the aluminium coverings, as observed in Figure 1.

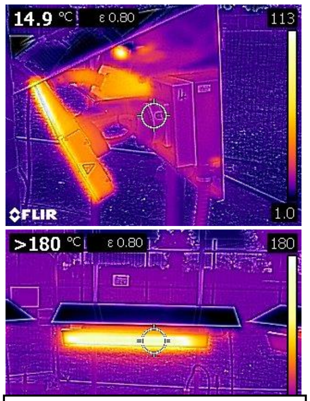

Figure 1. Thermal camera pictures Herschel heater fully ON.

2) **Two people are needed** for this step. Each heater and its covering weight ~15kg. Please use nitrile foam gloves in this step to prevent cuts (the sides of the aluminium coverings are sharp) and burns (the gloves are heatproof up to 149°C). One person holds the aluminium covering in the middle, from outside the plot (Figure 2, left picture). The other person unscrews the pieces that attaches the heater to the poll with the Allen key (Figure 2, right). The whole surface is not too heavy, one person can hold it, plus it won’t come down quickly as it is being partially supported by the attachment piece. Once both sides are unscrewed, each person holds the aluminium covering from each of the sides. Pull it up or down slowly from both sides at the same time.

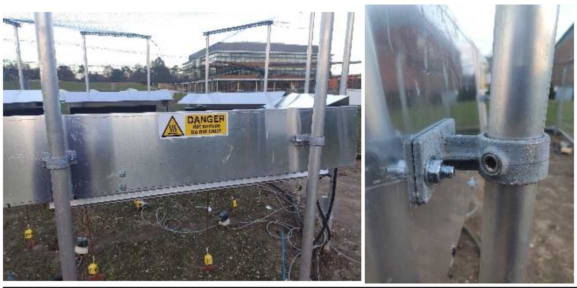

Figure 2. On the left, view of aluminium covering attached to the 2 polls. On the right, closer view of the attachment piece that holds the aluminium covering to the poll.

This is a controlled document maintained on electronic media. When appearing in paper form it should be checked against the master on the QA database to ensure that the current version is being used. This copy was printed on 16 February 2023.

3) Once the required height is reached, one of the two goes back to the middle to hold it while the other person screws both attachments back again.

### 4 WASTE DISPOSAL ARRANGEMENTS

N/A

### 5 EMERGENCY PROCEDURES

In an emergency dial 333 for assistance and a first aider. If there is no response and you need an ambulance dial 999/112.

### 6 RISK STATEMENT

All individuals using this procedure will be shown the risk assessment and given appropriate information, instruction and training in the risks and precautions necessary, including the use of any personal protective equipment required.

| ***SOP HEALTH RISK ASSESSMENT*** | | | |
|---|---|---|---|
| **[1] Activity:** | Adjusting the height of heaters in the free-air heating arrays (T-FACE system: Temperature Free-Air Controlled Enhancement) | | |
| **[2] Location of activity:** | Field station | | |
| **[3] Who is involved:** | Cristobal Uauy's team | | |
| **[4] Frequency of activity:** | Once per month | | |
| **[5] Duration of Activity:** | 40 mins | | |
| **[6] Chemical Hazard Name:** | **Hazard Statements** | **Route of Exposure*** | **Quantity Used** |
| N/A | | | |
| **[7] Details of biological agents of risk to human health: NA** | | | **GMRA Number** |
| **[8] Other Hazards:** | Please ■ as necessary | | |

This is a controlled document maintained on electronic media. When appearing in paper form it should be checked against the master on the QA database to ensure that the current version is being used. This copy was printed on 16 February 2023.

| | | | | | |
|---|---|---|---|---|---|
| Hot or Cold Burns | ☒ | Ionising Radiation** | ☐ | Ultra Violet or Infra Red | ☐ |
| Dust | ☐ | Noise | ☐ | Pollen Sensitizer | ☐ |
| Repetitive Action | ☐ | Extreme Cold Environment (< 0ºC) | ☐ | Lifting / Manual Handling | ☒ |
| Asphyxiation | ☐ | Cuts | ☒ | Electrical or electromagnetic fields | ☐ |
| Slips / trips / falls | ☒ | Display Screen Equipment | ☐ | Nanomaterials (man-made particles, tubes, rods, or fibres ≤100nm in size) | ☐ |
| Workplace Transport | ☐ | Lone Working or Remote Working | ☐ | Other (give details) | ☐ |

**[9] Control Measures:** Please ☒ as necessary

| | | | | | |
|---|---|---|---|---|---|
| Fume Cupboard | ☐ | Microbiological Safety Cabinet | ☐ | Total Containment Cabinet | ☐ |
| Ventilated Bench | ☐ | Spill Tray | ☐ | Trained personnel only | ☒ |
| Signs | ☐ | Reduce frequency/alternate activity | ☐ | Reduce duration of activity | ☐ |
| Sub divide a load | ☐ | 2 person lift of loads | ☒ | Not for more than 1 hour | ☐ |
| Regular, short breaks | ☐ | Alternate activities | ☐ | Local Exhaust Ventilation | ☐ |
| Screen / barrier | ☐ | Permit to work | ☐ | Other (give details) | ☐ |

**[10] Personal Protection:** Please ☒ as necessary

| | | | | | |
|---|---|---|---|---|---|
| Lab coat | ☐ | Safety Glasses | ☐ | Face Shield | ☐ |
| Goggles | ☐ | Gloves | ☒ | Thermal Protective Gloves | ☒ |
| Ear defenders | ☐ | Face fit tested face mask | ☐ | Airflow hood | ☐ |

This is a controlled document maintained on electronic media. When appearing in paper form it should be checked against the master on the QA database to ensure that the current version is being used. This copy was printed on 16 February 2023.

| | | | | | |
|---|---|---|---|---|---|
| Other (give details) | ☐ | | | | |
| **[11] Is personal monitoring and/or health surveillance required?** | | Yes ☐ | | No | ☒ |
| Details: | All new starters working in the laboratories should have a base level lung function assessment and skin surveillance assessment with Occupational Health. | | | | |
| **[12] Restrictions:** | | Please ☒ as necessary | | | |
| No lone working | ☐ | Not to be left unattended | ☐ | Named persons only | ☒ |
| In restricted area | ☐ | Not by pregnant or breastfeeding workers | ☐ | Under constant supervision | ☐ |
| Not by under 18's | ☒ | Other (give details) | ☐ | | |
| **[13] Level of Residual Risk**: | | Please ☒ as necessary | | | |
| Low | ☒ | Medium | ☐ | High | ☐ |
| Name of Assessor: Isabel Faci-Gomez | | | | Date: | 13/01/2023 |

* Route of exposure; S = skin, I = ingestion, B = inhalation

### 7 DOCUMENTATION

Add links to relevant H&S information on intranet or internet

Add any reference for any relevant manuals

Link to: Good Laboratory Practice in the Use of Chemicals:
http://intranet/cms/2199

Link to Biological and GM Safety:
http://intranet/cms/969

Link to Laboratory Waste Disposal:
http://intranet/cms/968

### 8 RELATED PROCEDURES

NONE

### 9 NOTES

The field-heated plots area has a restricted access. Please be careful as there are cables and electricity. Try not to touch anything that is not needed.

The wires connected to the heaters are shown in Figure 3 and the control panel in Figure 4.

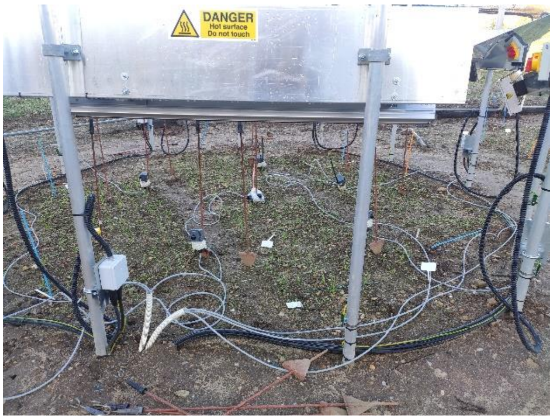

Figure 3. Waterproofed covered wires feeding heaters (on the top)

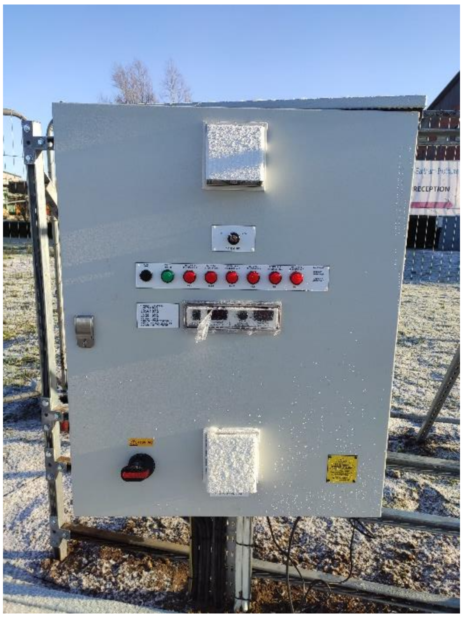

Figure 4. Control panel for the T-FACE system

### 10 APPENDICES

NONE

This is a controlled document maintained on electronic media. When appearing in paper form it should be checked against the master on the QA database to ensure that the current version is being used. This copy was printed on 16 February 2023.
